# An unbiased cell-culture selection yields DNA aptamers as novel senescent cell-specific reagents

**DOI:** 10.1101/2025.03.19.643361

**Authors:** Keenan S. Pearson, Sarah K. Jachim, Caroline D. Doherty, Brandon A. Wilbanks, Luis I. Prieto, Maria Dugan, Darren J. Baker, Nathan K. LeBrasseur, L. James Maher

**Affiliations:** Department of Biochemistry and Molecular Biology, Mayo Clinic College of Medicine and Science, Rochester, Minnesota 55905, United States; Department of Molecular Pharmacology and Experimental Therapeutics, Mayo Clinic College of Medicine and Science, Rochester, Minnesota 55905, United States; Department of Pediatric and Adolescent Medicine, Mayo Clinic, Rochester, Minnesota 55905, United States; Paul F. Glenn Center for Biology of Aging Research at Mayo Clinic, Mayo Clinic, Rochester, Minnesota 55905, United States; The Robert and Arlene Kogod Center on Aging, Mayo Clinic, Rochester, Minnesota 55905, United States; Department of Physical Medicine and Rehabilitation, Mayo Clinic, Rochester, Minnesota 55905, United States

## Abstract

Cellular senescence is an irreversible form of cell-cycle arrest caused by excessive stress or damage. While various biomarkers of cellular senescence have been proposed, there are currently no universal, stand-alone indicators of this condition. The field largely relies on the combined detection of multiple biomarkers to differentiate senescent cells from non-senescent cells. Here we introduce a new approach: unbiased cell culture selections to identify senescent cell-specific folded DNA aptamers from vast libraries of trillions of random 80-mer DNAs. Senescent mouse adult fibroblasts and their non-senescent counterparts were employed for selection. We demonstrate aptamer specificity for senescent mouse cells in culture, identify a form of fibronectin as the molecular target of two selected aptamers, show increased aptamer staining in naturally aged mouse tissues, and demonstrate decreased aptamer staining when p16 expressing cells are removed in a transgenic *INK-ATTAC* mouse model. This work demonstrates the value of unbiased cell-based selections to identify new senescence-specific DNA reagents.

## INTRODUCTION

Cellular senescence is a state of irreversible cell cycle arrest, first characterized by Hayflick and Moorhead in 1961.^1^ Since that time there has been extensive research to characterize mechanisms of senescence induction, the role of senescence in tissue remodeling, its anti-tumorigenesis function, its contribution to age-related diseases, and biomarkers to more specifically define senescent cells.^2-4^ While many proposed markers of senescence have been characterized, a single universal marker has not been defined. Combinations of various markers are generally needed to identify senescent cells. Common indicators include markers of cell cycle arrest, expression of cyclin dependent kinase inhibitors (CDKIs) p16 and p21,^5-8^ expressed features of the senescence-associated secretory phenotype (SASP) including cytokines such as IL6, chemokines such as MCP-1, and metalloproteinases,^9^ increased senescence-associated *β*-galactosidase (SA-*β*-gal) activity,^10^ and various morphological alterations such as cell size and shape.^11^ The accumulation of senescent cells in aging or in response to chemotherapeutic damage causes chronic inflammation due to the SASP and leads to increased pathogenesis.^8, 12^

Senolytics are a class of drugs intended to selectively clear senescent cells. These drugs must therefore selectively target senescent cells and avoid harm to quiescent or healthy, postmitotic cells. To improve targeting of senolytics, several modifications have been developed, including antibody-drug conjugates^13^ and lysosomal-dependent prodrugs.^14^ A similar strategy has been employed using an aptamer-functionalized liposome to deliver senolytics to a specific cell type, synoviocytes, associated with osteoarthritis.^15^ Recently, a strategy for highly specific targeting of a subpopulation of senescent cells was reported, which employed a three-part strategy. An aptamer targeting a membrane protein upregulated on the cells of interest was conjugated to a senolytic intended to selectively eliminate senescent cells by a self-immolating linker cleavable by SA-*β*-gal.^16^

Thus, these approaches attempt to identify markers upregulated in senescent cells of interest and modify a targeting moiety with the goal of increasing selective drug delivery. With such applications in mind, here we apply Systematic Evolution of Ligands by EXponential enrichment (SELEX) to identify DNA aptamers that distinguish carefully validated senescent cells from matched, healthy cells. We characterize the binding of several candidate aptamers using various cell lines in vitro. The molecular target of two of these aptamers is identified as a form of fibronectin. Binding activity of one of the identified aptamers is shown to exhibit a correlation with age and senescence burden in vivo.

## RESULTS

### Aptamer selection

Primary mouse ear fibroblasts were isolated from adult C57BL6/J mice (henceforth termed Mouse Adult Fibroblasts, MAFs). Senescence was induced in these cells using an etoposide regimen, and the status of the MAF cultures was confirmed by change in morphology, increased senescence associated β-galactosidase (SA-β-gal) activity, and upregulation of cyclin-dependent kinase inhibitors and SASP factors (Figure 1). Unchallenged MAFs were used as negative selection “control” targets and the etoposide-challenged senescent MAFs served as positive selection targets (Figure 2A). Selection progress over multiple rounds was monitored by subjecting recovered aptamer libraries to qPCR.^17^ A notable increase in library recovery was observed by round 7, and this increase was sustained through two additional rounds of selection (Figure 2B). To further assess the ability of late round aptamer library DNA to bind preferentially to senescent cells, the cell binding of samples of round 2 and round 8 libraries was compared. These aptamer libraries were exposed to senescent MAFs only, washed, recovered, and residual binding assessed by qPCR (Figure S1A). A significantly higher proportion of the round 8 library was recovered compared with the round 2 library (Figure S1B).

**Figure 1.**
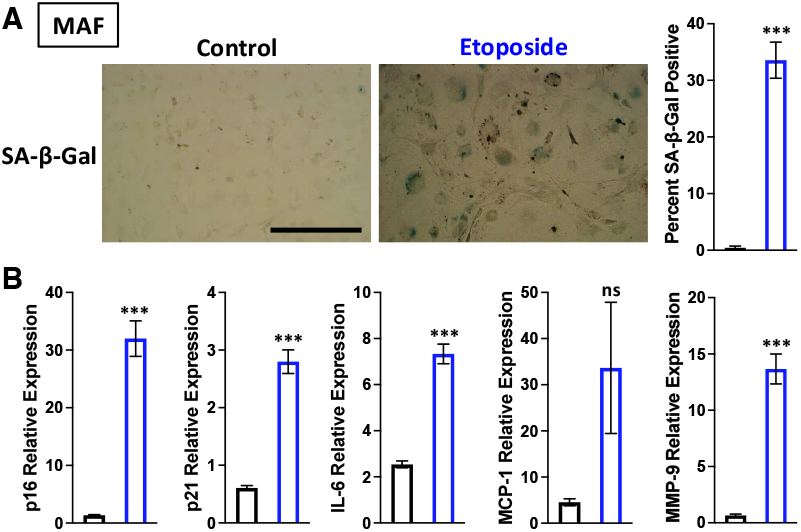
Validation of senescent phenotype in etoposide-challenged MAFs. A) Bright field images of control and etoposide-challenged MAFs show morphological changes and SA-β-Gal staining. The percent SA-β-Gal positive cells is quantified per field and compared using a two-tailed unpaired t-test (n=5; *p<0.05, **p<0.01, ***p<0.001). Scale bar is 400 µm. B) RT-qPCR mRNA quantitation of various markers of senescence assessed by ΔΔCT method normalized to TATAbox binding protein (TBP) mRNA as reference. Each comparison is made using a two-tailed unpaired t-test (n=6; *p<0.05, **p<0.01, ***p<0.001).

**Figure 2.**
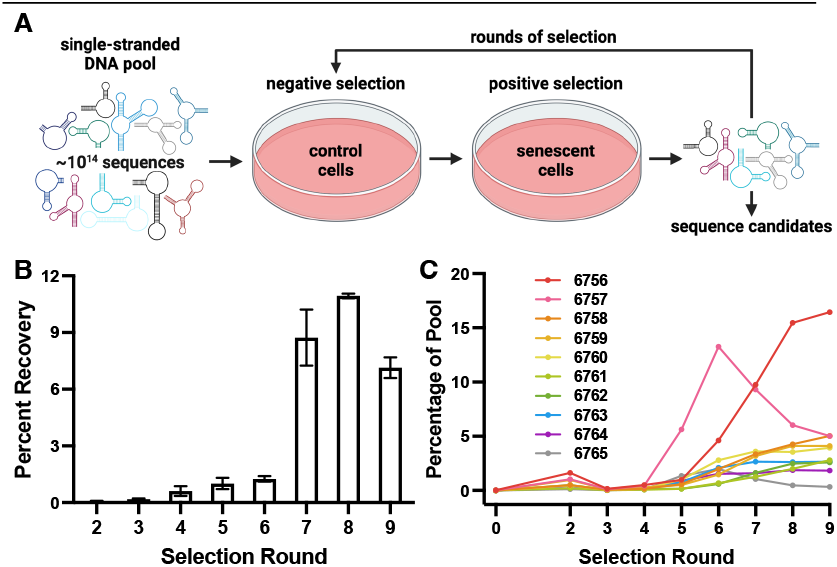
Selection of DNA aptamers that preferentially bind senescent cells. A) Schematic of selection procedure. B) qPCR quantification of library recovery across selection round. Error bars show standard error of technical replicates. C) Prevalence of selected aptamer candidates in deep sequencing data across rounds of selection.

DNA aptamers from all selection rounds and the original naïve library were subjected to Next Generation Sequencing to determine candidate aptamer sequences contributing to increased recovery observed by qPCR. The fraction of the libraries consisting of non-unique sequences substantially increased at round 5 and continued to increase through round 9 of selection, indicating an enrichment of senescent cell specific DNA molecules (Figure S1C). We synthesized ten unrelated 80-mer aptamer sequences and screened them for the ability to specifically bind to senescent cells (Figure 2C, Table S1). We also chose two negative control sequences present in the naïve library but not in any of the sequences recovered over 9 selection rounds (negative control oligonucleotides 6766 and 6767, Table S1).

### In vitro senescent cell binding screen

We assessed binding of the candidate sequences and controls to senescent and control MAFs by using qPCR to monitor recovery after incubation with cells, stringent washing, and cell lysis. These results indicated that all candidate aptamers, but not negative controls, bound to senescent cells more strongly than control cells (Figure 3A, Table S2). All aptamer candidates except 6764 exhibited statistically significant increased senescent cell binding relative to negative control oligonucleotide 6766 (Figure 3A). We then monitored cell binding of biotin-labeled candidates and controls using fluorescent streptavidin and quantified total fluorescence. Six of the candidates exhibited statistically significantly increased staining of senescent cells compared to 6766 (Figure 3B-C).

**Figure 3.**
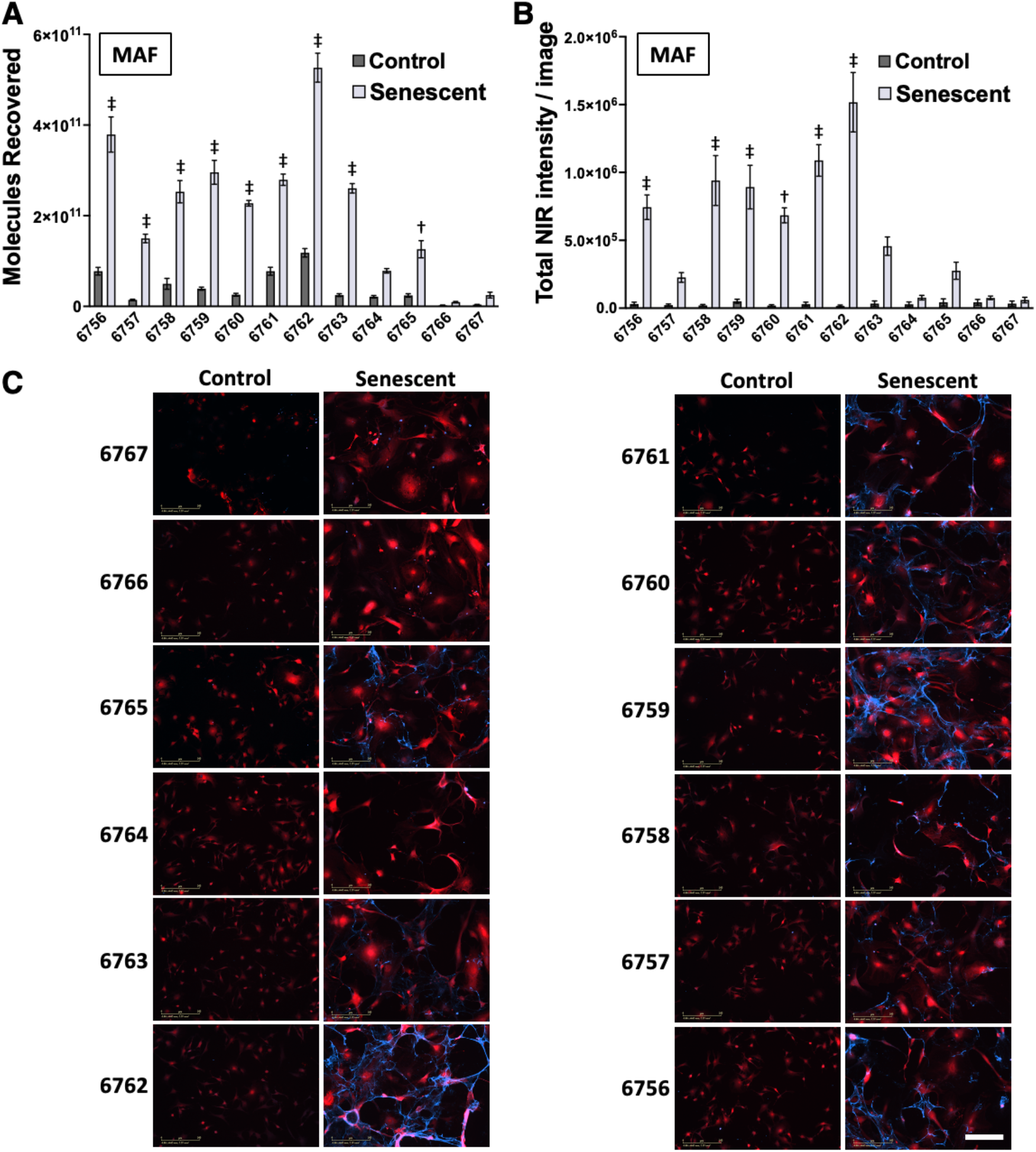
Candidate aptamer binding to senescent MAFs. A) Quantification of candidates (6756 – 6765) and controls (6766 and 6767) binding to control and senescent MAFs by qPCR. Aptamer concentration was 50 nM. Statistical significance is shown for candidates compared with control 6766 by one-way ANOVA with Dunnett’s post-hoc test for multiple comparisons (†: p<0.005, ‡: p<0.0005). Error bars indicate standard error for biological replicates (n=4). B) Quantification of staining by 50 nM aptamer/control calculated as total NIR signal intensity per image field. Statistical significance is shown for candidates compared with control 6766 by one-way ANOVA with Dunnett’s post-hoc test for multiple comparisons (*: p<0.05, †: p<0.005, ‡: p<0.0005). Error bars are shown as standard error for multiple image fields (n=9). C) Images taken with IncuCyte SX5 are shown at 20× magnification. Constitutively expressed TdTomato is shown in red, and aptamer staining is shown in blue (secondary stain with AlexaFluor647 labeled streptavidin binding biotinylated candidates and controls). Scale bar (lower right panel) is 200 µm.

To assess the importance of the different culture times prior to aptamer binding for senescent cells (8 d) vs. control cells (1 d), the growth of control MAFs was slowed by temporary serum starvation. An imaging experiment was repeated to assess aptamer binding under these conditions. Seven of the aptamer candidates exhibited statistically significantly increased staining relative to negative control 6766 (Figure S2A). In contrast, when senescent MAFs were treated with trypsin and replated the day before imaging, only two aptamers exhibited statistically significant staining relative to negative control 6766 (Figure S2B). These data suggest that senescent cell-specific aptamer binding is not an artifact due to prolonged senescent cell culture, but the data also suggest that the molecular target(s) of the selected aptamers are damaged by trypsin treatment or removed in the process of replating senescent MAFs.

Aptamer specificity was tested by subjecting MAFs to two alternative methods for senescence induction. X-ray irradiation has been a standard for in vitro senescence induction, and we again demonstrated typical senescence markers after using this methodology (Figure S3). All tested aptamers except 6764 exhibited statistically significant binding to X-ray induced senescent MAFs compared with the negative control oligonucleotide (Figure S4). Hydrogen peroxide exposure was also tested as a method for senescence induction. The resulting senescence phenotype was detectable, though less pronounced as determined by senescent marker characterization, therefore serving as a model of oxidative stress induced senescence (Figure S5). Again, all the aptamers except 6764 exhibited statistically significant binding to hydrogen peroxide-induced senescent MAFs compared with the negative control oligonucleotide (Figure S6). This demonstrates that the specificity of these aptamers for senescent cells is not changed by the induction method.

We tested the performance of candidate aptamers with a different senescent cell type. Mouse C2C12 myoblasts were challenged with etoposide and conventional markers of senescence were assessed (Figure S7). Five of the aptamers showed significant staining of senescent C2C12 cells compared to a negative control oligonucleotide (Figure S8). These five aptamers also demonstrated specific staining of senescent C2C12 cells compared to replicating control C2C12 cells (Table S2).

For further comparison, Normal Human Lung Fibroblasts (NHLF) and human IMR90 cells, were challenged with etoposide and the same increase in senescent markers was documented (Figures S9 and S10). Interestingly, none of the selected aptamers bound to the senescent NHLF or IMR90 cells above the level of the negative control oligonucleotide (Figure S11). These results demonstrate that the selected aptamers display binding specificity for senescent mouse cells (MAFs, C2C12) compared to senescent human cells (NHLF, IMR90).

### Binding target identification and characterization

To identify possible molecular targets, aptamers 6756 and 6762 were chosen for analysis. A SILAC-based proteomic assay was performed as described in Materials and Methods. Briefly, potential target proteins on isotopically labeled cultured cells were cross-linked to the tested biotinylated aptamer (6756 or 6762) using formaldehyde. A biotinylated negative control oligonucleotide, 6766, was similarly cross-linked but to non-isotopically labeled cells. Cell lysates were applied to streptavidin magnetic beads followed by stringent washing. Proteins were recovered by cross-link reversal at elevated temperatures. Heavy and light isotope labeled samples were combined as aptamer and negative control. Experiments were repeated with heavy and light isotope labeling reversed. After trypsin digestion, samples were subjected to LC-MS/MS analysis and peptide mass fingerprinting to identify proteins uniquely cross-linked to specific aptamers. Notably, peptides from only a single protein, fibronectin, were strongly crosslinked by both aptamers 6756 and 6762, displaying a selection ratio >1 for duplicate forward (heavy + aptamer/light + control) and duplicate reverse (light + aptamer/heavy + control) experiments (Table 1).

**Table 1.**
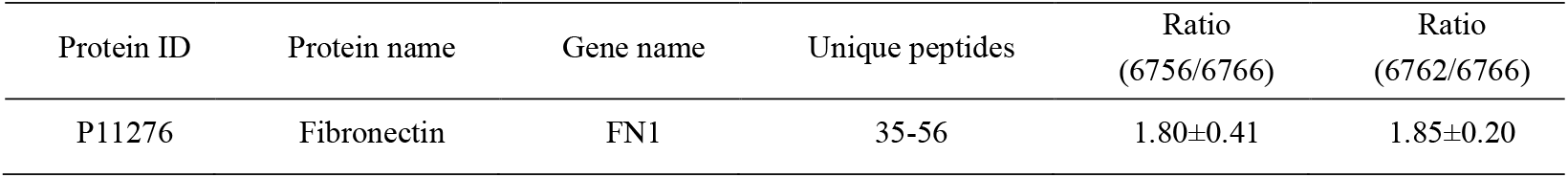
Results from SILAC based proteomics experiment show only fibronectin enriched in either 6756 or 6762 compared with negative control 6766.

To validate this nominated molecular binding partner, the same cross-link and pull-down approach was applied, followed by a Western blot. This experiment showed that aptamers 6756 and 6762 selectively enriched fibronectin compared to negative control oligonucleotide 6766 or streptavidin magnetic beads only (Figures 4A and S12). Further, purified native mouse fibronectin was exposed to immobilized aptamer or negative control oligonucleotide displayed on streptavidin magnetic beads without formaldehyde cross-linking. This procedure resulted in fibronectin capture by aptamers 6756 and 6762 when compared with negative control oligonucleotide 6766 or beads alone (Figure 4B).

**Figure 4.**
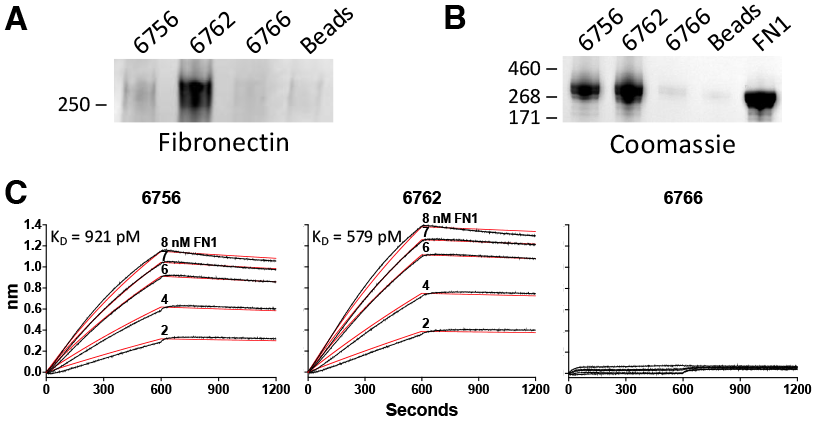
Identification and verification of target protein of aptamers 6756 and 6762. A) Aptamer pull-down from cell lysates followed by SDS-PAGE and Western blotting show fibronectin to be enriched in both 6756 and 6762 compared to negative controls. B) Purified fibronectin from mouse plasma was incubated with aptamer or negative control loaded streptavidin magnetic beads. Beads were washed and any remaining protein was eluted and analyzed by SDS-PAGE. The gel was stained with Coomassie blue and imaged. C) Biolayer Interferometry result shows dissociation constants of 6756 (921 pM) and 6762 (579 pM), and no binding of negative control 6766, with purified fibronectin from mouse plasma.

The affinity of 6756 and 6762 for fibronectin was assessed by biolayer interferometry (BLI). Aptamers 6756 and 6762 exhibited equilibrium dissociation constants of 921 pM and 579 pM respectively (Figure 4C). The negative control oligonucleotide 6766 displayed no appreciable binding (Figure 4C).

Confocal microscopy was used to assess colocalization of anti-fibronectin antibody staining and staining by aptamers 6756 or 6762 on cultured cells. Interestingly, the staining patterns are subtly different, suggesting that a variant form of fibronectin may be detected by senescent cell-specific aptamers (Figure S13). Confocal microscopy was also utilized to assess the relative amount of anti-fibronectin antibody staining on control and senescent mouse C2C12 cells and human IMR90 cells. Both cell lines showed an increased amount of fibronectin staining in senescent cells versus control cells. There was not a statistically significant difference in the amount of fibronectin staining between senescent C2C12 and senescent IMR90 cells (Figure S14).

Levels of *Fn1* mRNA transcripts (encoding fibronectin) from RNA-seq data were compared between etoposide-induced senescent MAFs and control MAFs. Intriguingly, *Fn1* transcript expression is not statistically different between the two in vitro conditions (Figure S15), implying senescent cells may have a different abundance, location, or isoform of fibronectin that is only detectable at the protein level.

### Aptamer stain correlation with in vivo age and senescence burden

To assess whether DNA aptamer specificity for senescent cells in vitro correlates with binding to tissue from animals of different ages, we compared senescence-specific aptamer 6762 to negative control 6766 for staining lung tissue sections from naturally aged mice. There is no appreciable staining in younger mice, and we observed that for ages 22 and 30 months there is an obvious increase in specific aptamer staining across the tissue sections that is much stronger than for the negative control (Figure 5). Higher magnification images show that this staining varies by region of the tissue (Figure S16). This increased staining with age is consistent with the known age-associated increase in senescent cell burden.

**Figure 5.**
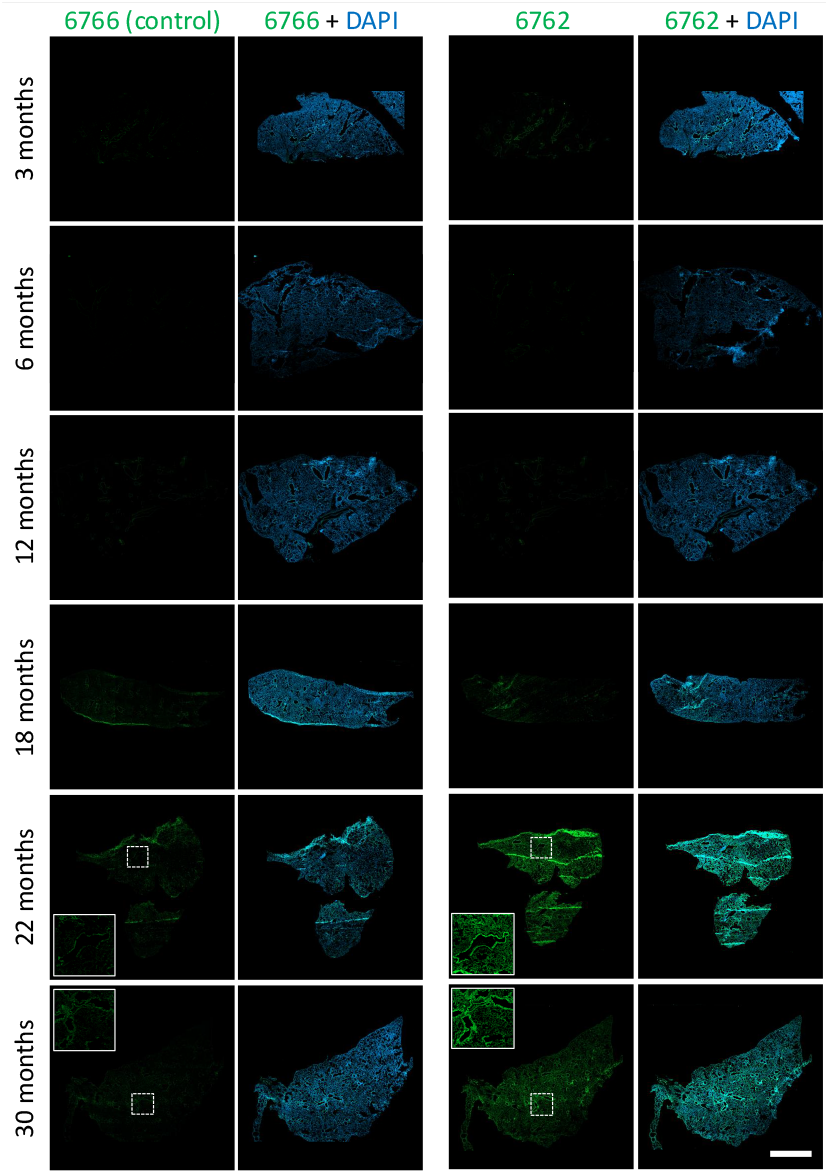
Aptamer 6762 staining of mouse lung tissue increases with age. Aptamer 6762 and negative control oligonucleotide 6766 are directly detected using the fluorescein label (green). Nuclei are stained with DAPI (blue). Inlays are 3×enlargements of indicated regions. The images are 5× tiled confocal images of the entire tissue section. Scale bar (lower right panel) is 2 mm.

To further assess whether age-related staining of tissue sections correlates with in vivo senescent cell burden, we stained lung tissues from 21-month-old *INK-ATTAC* mice either treated with AP (selectively eliminates senescent cells^18^) or with vehicle. AP-treated tissues showed a statistically significant decrease in 6762 staining compared with vehicle (Figures 6, S17, and S18). In this same experiment, there was no change in anti-fibronectin antibody staining (Figure 6, S17, and S18). Interestingly, higher magnification (40×) images show that, as in the in vitro experiment, anti-fibronectin antibody and 6762 stains do not show the same pattern of target interaction (Figure S19).

**Figure 6.**
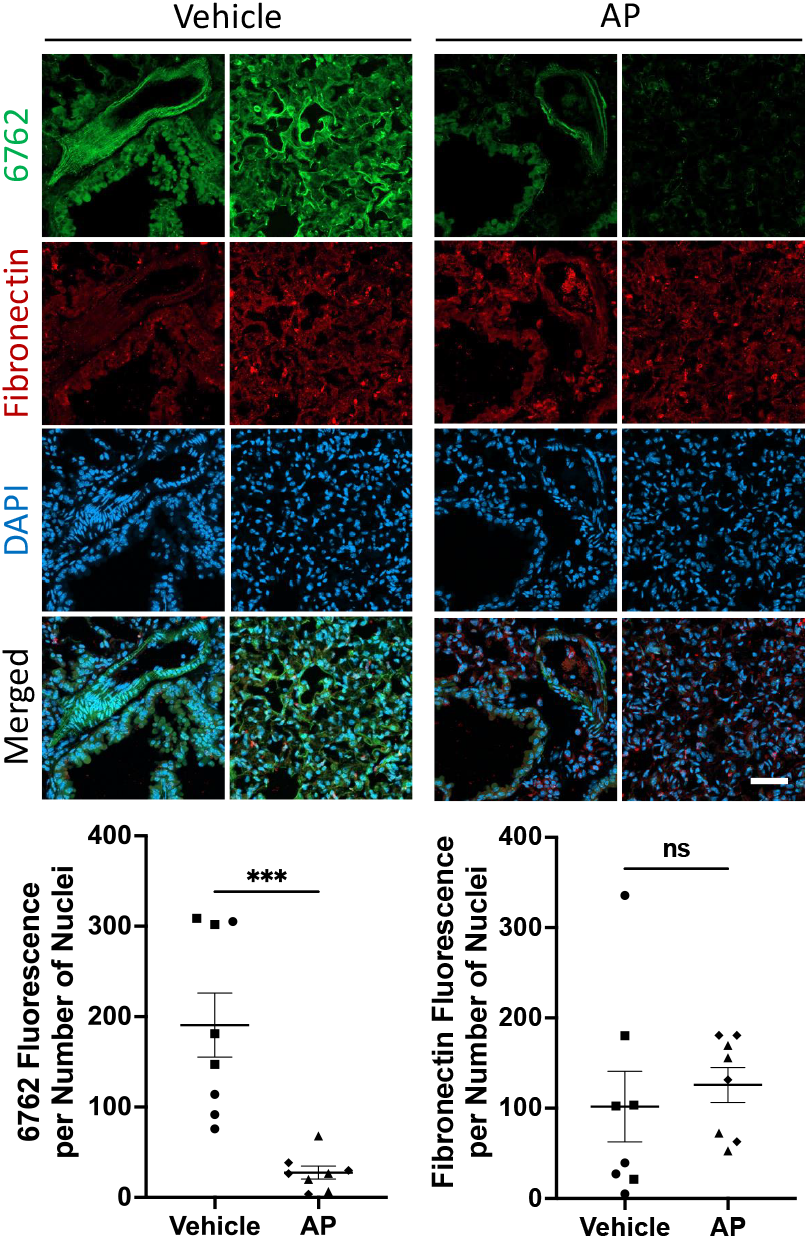
Removal of p16-positive cells leads to reduced aptamer 6762 (but not fibronectin) staining. Aptamer 6762 is detected by fluorescein label (green). Fibronectin antibody is detected with AlexaFluor594 anti-rabbit secondary (red). Nuclei are stained with DAPI (blue). Quantification of aptamer 6762 fluorescence per nuclei and fibronectin fluorescence per nuclei was obtained from 8 image fields per condition. Each comparison is made using a two-tailed unpaired t-test (***p<0.001, ns: p>0.05). Scale bar (lower right panel) is 50 µm.

## DISCUSSION

The results reported here suggest the successful identification of a collection of DNA aptamers with varying specificities for senescent cells in culture. These results extend to various cell types (fibroblasts and myoblasts) and various methods for senescence induction (etoposide challenge, X-ray irradiation, and hydrogen peroxide exposure). Our results further suggest that the selected aptamers demonstrate binding preference for senescent mouse cells over senescent human cells. This selectivity might be explored by screening additional cell lines.

The apparent fibronectin specificity of lead selected aptamers is interesting considering the mouse vs. human selectivity and the evident difference in cell staining pattern by selected aptamers and conventional anti-fibronectin antibodies. It may be possible to review epitope differences and posttranslational modifications that vary between mouse and human fibronectin to understand these results. Because the staining patterns between the anti-fibronectin antibody and aptamers 6756 (in vitro) and 6762 (in vitro and in vivo) do not match, the aptamer target epitope may represent a unique fibronectin form produced during senescence. This concept is further supported by our results showing that fibronectin expression is constant in senescent mouse and senescent human cells in culture as judged by an anti-fibronectin antibody.

It has been previously argued that fibronectin protein levels increase in cell culture models of senescence.^19^ While we do not detect increases in fibronectin mRNA upon induction of senescence, we do show an increase in staining with an anti-fibronectin antibody. However, this did not hold true in tissue staining. We interpret our results to show that some unique senescence-specific form of fibronectin is detected during the selection of multiple aptamers in our unbiased cell culture selection. It will be of future interest to determine what fibronectin variant is detected by the aptamers we have selected and if this variant can be of value as a biomarker, potential diagnostic/prognostic marker, and possible target for homing senolytics. We again note that if fibronectin protein levels increase with senescence, this is not accompanied by an increase in fibronectin mRNA.^20^

Our work further demonstrates that binding of representative selected aptamer 6762 is dependent on age and senescent cell burden in tissue sections. In contrast, binding of the anti-fibronectin antibody we studied did not correlate with senescence burden. These results again support the notion that 6762 recognizes a fibronectin epitope uniquely present in senescence. Alternatively, a component of aptamer selectivity vs. anti-fibronectin antibodies could be due to the smaller size of DNA aptamers and corresponding greater tissue section penetration. However, this would not explain the difference in species specificity between the reagents.

Aptamer 6762 stains widely across all mouse tissue sections associated with advanced age and senescent cell burden. If 6762 is truly selective for senescent cells, this staining result would indicate a much higher prevalence of senescent cells than previous work has shown in animals at advanced age.^21^ Because 6762 binds some form of fibronectin, a component of the extracellular matrix (ECM), we hypothesize that some of the observed staining may not directly indicate the presence of senescent cells, but perhaps ECM residue left behind by senescent cells that have migrated or been cleared over time. Additionally, the powerful paracrine signaling from the SASP of senescent cells can remodel ECM and change the local tissue environment, even when senescent cells are rare. Together, these observations highlight an age-associated change in fibronectin in tissues. This will be an important consideration moving forward in targeting age-related or senescent-related targets in vivo.

It is interesting to speculate as to the exact fibronectin specificity of 6756 and 6762, given that cellular staining does not precisely match that of conventional anti-fibronectin antibodies (e.g. Figure S13). One possibility is that 6756 and/or 6762 have specificity for fibronectin containing Extra Domain A (EDA), a unique splice variant that has been shown to be more highly expressed in tissues with higher senescence burden.^22^ If 6756 or 6762 preferentially binds EDA but also cross-reacts to some extent with related Type III fibronectin domains, this could help to explain the observed staining patterns. In support of this hypothetical anti-fibronectin-EDA aptamer specificity, we note that EDA peptides were detected among those recovered after 6756 and 6762 cross-linking to senescent cells.

It is possible to envision future aptamer selections to identify aptamers for senescent human cells. Selections might also be designed to identify aptamers that are internalized into senescent cells. Such tools might eventually be applied prognostically to identify individuals most likely to benefit from senolytic treatment. Similar aptamers might be applied in the development of aptamer-drug conjugates that selectively deliver drugs to senescent cells. Such approaches might improve the specificity of senolytics and senomorphics.

The complexity of the senescence phenotype and the lack of a single universal biomarker to reliably identify senescent cells have posed significant challenges in the field of aging research. In this study, we leveraged an unbiased cell-based selection technique and demonstrated its value in identifying new senescence-specific DNA reagents.

## ASSOCIATED CONTENT

### Supporting Information

The Supporting Information is available free of charge on the ACS Publications website.

Materials and Methods, supplementary tables, and supplementary figures (PDF)

## AUTHOR INFORMATION

### Notes

DJB has a potential financial interest related to this research. He is a co-inventor on patents held by Mayo Clinic, patent applications licensed to or filed by Unity Biotechnology, and a Unity Biotechnology shareholder. Research in the Baker laboratory has been reviewed by the Mayo Clinic Conflict of Interest Review Board and is being conducted in compliance with Mayo Clinic Conflict of Interest policies. The other authors declare no competing interests.

## ACKNOWLEDGMENT

The authors thank members of the Maher, LeBrasseur, and Baker laboratories. We acknowledge the Mayo Clinic Genome Analysis Core for their contributions. We thank the staff of the Mayo Clinic Proteomics Core, which is a shared resource of the Mayo Clinic Cancer Center (NCI P30 CA15083).

## ABBREVIATIONS

SELEX: systematic evolution of ligands by exponential enrichment
SASP: senescence-associated secretory phenotype
MAF: mouse adult fibroblast.

## Graphical TOC

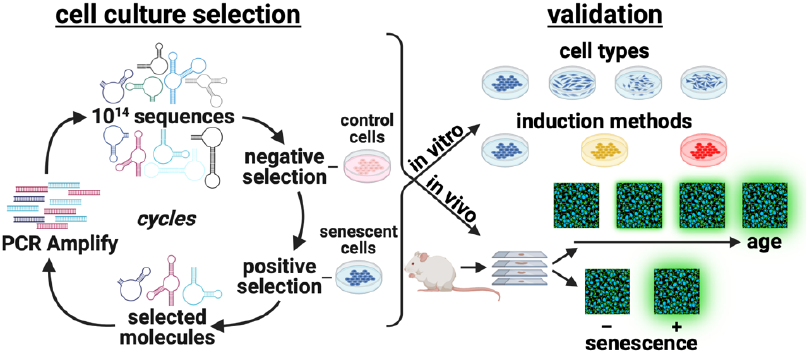

## REFERENCES

(1) Hayflick, L.; Moorhead, P. S. The serial cultivation of human diploid cell strains. Exp Cell Res 1961, 25, 585–621. DOI: 10.1016/0014-4827(61)90192-6 From NLM Medline.

(2) Dodig, S.; Cepelak, I.; Pavic, I. Hallmarks of senescence and aging. Biochem Med (Zagreb) 2019, 29 (3), 030501. DOI: 10.11613/BM.2019.030501 From NLM Medline.

(3) Hernandez-Segura, A.; Nehme, J.; Demaria, M. Hallmarks of Cellular Senescence. Trends Cell Biol 2018, 28 (6), 436–453. DOI: 10.1016/j.tcb.2018.02.001 From NLM Medline.

(4) Munoz-Espin, D.; Serrano, M. Cellular senescence: from physiology to pathology. Nat Rev Mol Cell Biol 2014, 15 (7), 482–496. DOI: 10.1038/nrm3823 From NLM Medline.

(5) Beausejour, C. M.; Krtolica, A.; Galimi, F.; Narita, M.; Lowe, S. W.; Yaswen, P.; Campisi, J. Reversal of human cellular senescence: roles of the p53 and p16 pathways. EMBO J 2003, 22 (16), 4212–4222. DOI: 10.1093/emboj/cdg417 From NLM Medline.

(6) Dulic, V.; Kaufmann, W. K.; Wilson, S. J.; Tlsty, T. D.; Lees, E.; Harper, J. W.; Elledge, S. J.; Reed, S. I. p53-dependent inhibition of cyclin-dependent kinase activities in human fibroblasts during radiation-induced G1 arrest. Cell 1994, 76 (6), 1013–1023. DOI: 10.1016/0092-8674(94)90379-4 From NLM Medline.

(7) el-Deiry, W. S.; Tokino, T.; Velculescu, V. E.; Levy, D. B.; Parsons, R.; Trent, J. M.; Lin, D.; Mercer, W. E.; Kinzler, K. W.; Vogelstein, B. WAF1, a potential mediator of p53 tumor suppression. Cell 1993, 75 (4), 817–825. DOI: 10.1016/0092-8674(93)90500-p From NLM Medline.

(8) Krishnamurthy, J.; Torrice, C.; Ramsey, M. R.; Kovalev, G. I.; Al-Regaiey, K.; Su, L.; Sharpless, N. E. Ink4a/Arf expression is a biomarker of aging. J Clin Invest 2004, 114 (9), 1299–1307. DOI: 10.1172/JCI22475 From NLM Medline.

(9) Coppe, J. P.; Desprez, P. Y.; Krtolica, A.; Campisi, J. The senescence-associated secretory phenotype: the dark side of tumor suppression. Annu Rev Pathol 2010, 5, 99–118. DOI: 10.1146/annurev-pathol-121808-102144 From NLM Medline.

(10) Dimri, G. P.; Lee, X.; Basile, G.; Acosta, M.; Scott, G.; Roskelley, C.; Medrano, E. E.; Linskens, M.; Rubelj, I.; Pereira-Smith, O.; et al. A biomarker that identifies senescent human cells in culture and in aging skin in vivo. Proc Natl Acad Sci U S A 1995, 92 (20), 9363–9367. DOI: 10.1073/pnas.92.20.9363 From NLM Medline.

(11) Bayreuther, K.; Rodemann, H. P.; Hommel, R.; Dittmann, K.; Albiez, M.; Francz, P. I. Human skin fibroblasts in vitro differentiate along a terminal cell lineage. Proc Natl Acad Sci U S A 1988, 85 (14), 5112–5116. DOI: 10.1073/pnas.85.14.5112 From NLM Medline.

(12) Campisi, J.; d’Adda di Fagagna, F. Cellular senescence: when bad things happen to good cells. Nat Rev Mol Cell Biol 2007, 8 (9), 729–740. DOI: 10.1038/nrm2233 From NLM Medline.

(13) Poblocka, M.; Bassey, A. L.; Smith, V. M.; Falcicchio, M.; Manso, A. S.; Althubiti, M.; Sheng, X.; Kyle, A.; Barber, R.; Frigerio, M.; et al. Targeted clearance of senescent cells using an antibody-drug conjugate against a specific membrane marker. Sci Rep 2021, 11 (1), 20358. DOI: 10.1038/s41598-021-99852-2 From NLM Medline.

(14) Cai, Y.; Zhou, H.; Zhu, Y.; Sun, Q.; Ji, Y.; Xue, A.; Wang, Y.; Chen, W.; Yu, X.; Wang, L.; et al. Elimination of senescent cells by beta-galactosidase-targeted prodrug attenuates inflammation and restores physical function in aged mice. Cell Res 2020, 30 (7), 574–589. DOI: 10.1038/s41422-020-0314-9 From NLM Medline.

(15) Chen, X.; Zhang, L.; Shao, X. Y.; Gong, W.; Shi, T. S.; Dong, J.; Shi, Y.; Shen, S. Y.; He, Y.; Qin, J. H.; et al. Specific Clearance of Senescent Synoviocytes Suppresses the Development of Osteoarthritis based on Aptamer-Functionalized Targeted Drug Delivery System. Adv Funct Mater 2022, 32 (17). DOI: ARTN 2109460 10.1002/adfm.202109460.

(16) Xia, Y.; Li, J.; Wang, L.; Xie, Y.; Zhang, L.; Han, X.; Tan, W.; Liu, Y. Engineering Hierarchical Recognition-Mediated Senolytics for Reliable Regulation of Cellular Senescence and Anti-Atherosclerosis Therapy. Angew Chem Int Ed Engl 2023, 62 (4), e202214169. DOI: 10.1002/anie.202214169 From NLM Medline.

(17) Pearson, K.; Doherty, C.; Zhang, D.; Becker, N. A.; Maher, L. J., 3rd. Optimized quantitative PCR analysis of random DNA aptamer libraries. Anal Biochem 2022, 650, 114712. DOI: 10.1016/j.ab.2022.114712 From NLM Medline.

(18) Baker, D. J.; Childs, B. G.; Durik, M.; Wijers, M. E.; Sieben, C. J.; Zhong, J.; Saltness, R. A.; Jeganathan, K. B.; Verzosa, G. C.; Pezeshki, A.; et al. Naturally occurring p16(Ink4a)-positive cells shorten healthy lifespan. Nature 2016, 530 (7589), 184–189. DOI: 10.1038/nature16932 From NLM Medline.

(19) Kumazaki, T.; Robetorye, R. S.; Robetorye, S. C.; Smith, J. R. Fibronectin expression increases during in vitro cellular senescence: correlation with increased cell area. Exp Cell Res 1991, 195 (1), 13–19. DOI: 10.1016/0014-4827(91)90494-f From NLM Medline.

(20) Casella, G.; Munk, R.; Kim, K. M.; Piao, Y.; De, S.; Abdelmohsen, K.; Gorospe, M. Transcriptome signature of cellular senescence. Nucleic Acids Res 2019, 47 (21), 11476. DOI: 10.1093/nar/gkz879 From NLM PubMed-not-MEDLINE.

(21) Biran, A.; Zada, L.; Abou Karam, P.; Vadai, E.; Roitman, L.; Ovadya, Y.; Porat, Z.; Krizhanovsky, V. Quantitative identification of senescent cells in aging and disease. Aging Cell 2017, 16 (4), 661–671. DOI: 10.1111/acel.12592 From NLM Medline.

(22) Sueblinvong, V.; Neveu, W. A.; Neujahr, D. C.; Mills, S. T.; Rojas, M.; Roman, J.; Guidot, D. M. Aging promotes pro-fibrotic matrix production and increases fibrocyte recruitment during acute lung injury. Adv Biosci Biotechnol 2014, 5 (1), 19–30. DOI: 10.4236/abb.2014.51004 From NLM PubMed-not-MEDLINE.

