## Supplementary data for "An unbiased cell-culture selection yields DNA aptamers as novel senescent cell-specific reagents"

**Table of Contents:**

**Materials and Methods**

**Supplementary Tables**

Table S1. Names and primary structures of primers, libraries, and aptamers.
Table S2. Statistical comparison of aptamer binding in control vs senescent conditions.

**Supplementary Figures**

Figure. S1. Verification of overall pool behavior and deep sequencing data.
Figure. S2. Effects of cell culture time vs. replating on aptamer binding.
Figure. S3. Validation of senescent phenotype in etoposide-treated C2C12 cells.
Figure. S4. Screening candidate aptamer binding to senescent C2C12 cells.
Figure. S5. Validation of senescent phenotype in etoposide-treated NHLF cells.
Figure. S6. Validation of senescent phenotype in etoposide-treated IMR90 cells.
Figure. S7. Screening candidate aptamer binding to senescent NHLF and IMR90 cells.
Figure. S8. Validation of senescent phenotype in X-ray irradiated MAFs.
Figure. S9. Screening candidate aptamer binding to X-ray-induced senescent MAFs.
Figure. S10. Validation of senescent phenotype in hydrogen peroxide treated MAFs.
Figure. S11. Screening candidate aptamer binding to hydrogen peroxide-induced senescent MAFs.
Figure. S12. Quality control data for anti-fibronectin Western blot.
Figure. S13. Confocal imaging of senescent C2C12 cells co-stained with aptamers and anti-fibronectin antibody do not show complete overlap.
Figure. S14. Fibronectin staining is elevated in both senescent C2C12 and IMR90 cells compared with control cells.
Figure. S15. RNA-seq relative quantitation of *FN1* transcripts.
Figure. S16. Aptamer 6762 staining of naturally aged mouse lung tissue varies by region.
Figure. S17. Removal of p16-positive cells leads to reduced aptamer 6762 staining throughout the entire tissue section.
Figure. S18. Additional images show removal of p16-positive cells leads to reduced aptamer 6762 staining.
Figure. S19. Confocal imaging of mouse lung tissue with aptamers and anti-fibronectin antibody do not show complete overlap.

**Supplementary References**

### Materials and Methods

#### Cell culture

Primary mouse ear fibroblasts were isolated from adult C57BL6/J mice as previously described with minimal modifications (1). Tissue digestion was performed with collagenase Type 2 (Worthington) at 800U/mL in DMEM with 1% penicillin-streptomycin-glutamine and 25 mM HEPES buffer (Gibco) for 1 h at 37°C. Cells and digested tissue were washed with HBSS and digested for an additional 20 min in 0.05% trypsin (Gibco) at 37°C. Isolated cells were then cultured in DMEM with 10% FBS and 1% penicillin-streptomycin-glutamine (Gibco) and provided fresh media every 3 d. These cells were maintained at 3% oxygen and 5% $CO_2$ and used in experiments between passage 3 and passage 8.

Mouse myoblasts (C2C12; ATCC), primary human lung fibroblasts (NHLF; Lonza), and human fibroblasts (IMR90; ATCC) were cultured DMEM with 10% FBS and 1% penicillin-streptomycin-glutamine (Gibco) and provided fresh media every 3 d. These cells were maintained at 20% oxygen and 5% $CO_2$. NHLF were used in experiments between passage 3 and passage 6.

Etoposide-induced senescent cells were exposed to 20 μM etoposide (Sigma) for 48 h, then cultured for 5 d. X-ray-induced senescent cells were exposed to 10 Gy radiation using an RS2000 X-Ray inducer (RAD Source Technologies), then cultured for 21 d. Oxidative damage-induced senescent cells were exposed to 400 μM $H_2O_2$ for 48 h in serum free media, then cultured for 10 d. To control for the time in culture, a serum starvation protocol was used in one experiment: the control MAFs were grown with 0% FBS for 5 d, 1% FBS for 2 d, and 5% FBS for 1 d.

**Senescence-associated β-galactosidase (SA-β-Gal) staining**

Cells were fixed in phosphate-buffered 4% paraformaldehyde for 10 min at room temperature. Cells were washed twice with PBS and incubated 16–18 h in the SA–β-Gal staining solution (1 mg/mL X-Gal, 40 mM citric acid/sodium phosphate buffer pH 6.0, 5 mM potassium ferrocyanide, 5 mM potassium ferricyanide, 150 mM sodium chloride, and 2 mM magnesium chloride) on a shaker and in the dark at 37°C. Cells were washed twice with PBS and stained with Hoechst dye for 10 min as a nuclear stain. Images were taken under bright field for SA–β-Gal staining and then the same field under blue fluorescence channel for nuclear staining (EVOS 5000).

**RNA isolation, cDNA synthesis, and qPCR**

RNA was isolated according to manufacturer's instructions using TRIzol reagent (Invitrogen). RNA concentration was assessed by Nanodrop 8000 (Thermo Fisher Scientific). cDNA was synthesized using M-MLV reverse transcriptase (Invitrogen) and real-time PCR was performed with PerfeCTa FastMix II (QuantaBio) and the Applied Biosystems StepOne Plus Real-Time PCR system (Applied Biosystems). Gene expression was analyzed by the ΔΔCT method and normalized to TATA-box binding protein (TBP) mRNA as a reference.

**Buffers**

Wash buffer is defined as 1× DPBS (Gibco, 14190-144) containing 5 mM $MgCl_2$ and 4.5 g/L glucose. Binding buffer is defined as 1× DPBS containing 5 mM $MgCl_2$, 4.5 g/L glucose,

100 μg/mL tRNA, and 100 μg/mL BSA. Lysis buffer is defined as 10 mM Tris-HCl, pH 7.5, 10 mM NaCl, and 0.5% IGEPAL. Elution buffer is defined as 10 mM Tris-HCl, pH 8.0, 10 mM EDTA, 5 mM DTT, and 1% SDS. Binding and washing (B&W) buffer (2×) for Streptavidin Dynabeads M-270 (Invitrogen, 65305) is defined as 10 mM Tris-HCl at pH 7.5, 1 mM EDTA, and 2 M NaCl.

**Oligonucleotides**

DNA library, primers, and aptamers were purchased from Integrated DNA Technologies (Coralville, IA) with standard desalting. All sequences and modification abbreviations are provided in Table S1.

**DNA aptamer selection**

Senescent MAFs were cultured as described above to ~80% confluency on untreated 10-cm tissue culture dishes (Falcon, 353003). Control MAFs were plated the day before selection and cultured as described above to ~80% confluency on untreated 10-cm tissue culture dishes. Naïve DNA library (6643) was purchased from Integrated DNA Technologies with standard desalting (Coralville, IA). For round 1, 1 nmol (~$6\times10^{14}$ molecules) random DNA library 6643 was prepared in 2 mL wash buffer, heated at 90 ˚C for 5 min, and snap cooled on ice. tRNA (100 μg/mL) and BSA (100 μg/mL) were then added as non-specific competitors. Subsequent rounds 2-9 were prepared using 100 pmol library purified after large-scale PCR amplification. Medium was aspirated from control MAFs, and the dish was rinsed with warm DPBS once. The library solution was added to the control MAFs and incubated at 37 °C for 30 min. Media was aspirated from senescent MAFs and the dish was rinsed with warm DPBS once. The library solution was transferred from the control MAFs to the senescent MAFs and incubated at 37 °C for 30 min. The senescent MAFs were washed on the dish three times with 2 mL wash buffer. The cells were scraped into wash buffer and collected by centrifugation (500×g for 5 min). The supernatant was removed, and the cell pellet was resuspended in 500 μL wash buffer. Cells were then lysed by heating at 95 °C for 10 min. Debris was removed by centrifugation at 13,100×g for 5 min, and the supernatant containing the recovered library was moved to a fresh tube.

In the first round only the cell pellet was resuspended in water instead and used in its entirety as the template for PCR with the following reagent volumes: 500 μL round 1 lysate, 100 μL 10× PCR buffer, 80 μL 2.5 mM dNTPs, 100 μL 1 mg/mL BSA, 80 μL 50 mM $MgCl_2$, 50 μL 6638 (10 μM stock), 50 μL 6644 (10 μM stock), 20 μL water, 20 μL Taq (Invitrogen, 10342178). The mixture was split into 100 μL aliquots and subjected to thermal cycling using the following protocol: 95 °C, 60s; 10×(94 °C, 30s; 50 °C, 35s; 72 °C, 30s).

The lysate (or the completed PCR solution from the first round) was used directly as a template for analytical PCR to determine the optimum number of cycles to be used in a large-scale PCR. The following reagent volumes were used: 200 μL template, 200 μL 10× PCR buffer, 160 μL 2.5 mM dNTPs, 200 μL 1 mg/mL BSA, 160 μL 50 mM $MgCl_2$, 200 μL 6638 (5 μM stock), 200 μL 6644 (5 μM stock), 652 μL water, 28 μL Taq. The mixture was split into 100 μL aliquots and subjected to thermal cycling following the protocol described above with the optimum number of cycles as determined with analytical PCR.

Following amplification, aliquots were recombined. 0.1 volume of 3M NaOAc was added and mixed. Nucleic acids were precipitated by the addition of 2.5 volumes of ethanol, with mixing, incubation on dry ice for 15 min, and centrifugation at 17,000×g for 15 min. The pellet was washed with 70% ethanol followed by centrifugation at 17,000×g for 5 min. The pellet was air dried, resuspended in 40 µL water, combined with 160 µL deionized formamide, heated at 90˚C for 5 min, and loaded onto a 10% denaturing polyacrylamide gel (7.5 M urea, 19:1 acrylamide:bisacrylamide) and subjected to electrophoresis for 2 h at 600 V (26.25 V/cm).

Bands were visualized with handheld UV lamp. The higher mobility DNA band, comprised of the fluorescent single stranded library, was excised using a clean razor blade. The band was cleaved into small cubes and eluted overnight at 37˚C in 500 µL 2× PK buffer (100 mM Tris-HCl (pH 7.5), 200 mM NaCl, 2 mM EDTA, 1% SDS) on an end-over-end rotator. The supernatant extracted with an equal volume of phenol:chloroform:iso-amyl alcohol (25:24:1) (VWR, 97064-692). The upper aqueous phase was then precipitated from ethanol as described above. The pellet was resuspended in 100 µL water. The concentration of the library was estimated using a molar extinction coefficient at 260 nm (766,875 $M^{-1}cm^{-1}$).

**Monitoring library recovery**

qPCR reactions (30 µL) were performed with the following reagent volumes: 15 µL PerfeCTa SYBR Green FastMix for IQ (Quantabio, 95071-012), 3 µL 5 µM forward primer, 3 µL 5 µM reverse primer, 7.5 µL water, 1.5 µL template. Samples were transferred to a CFX96 Touch Deep Well Real-Time PCR Detection System (Bio-Rad Laboratories, 1854095) and subjected to thermal cycling using a protocol of 95 ˚C, 30s; 40×(95 ˚C, 15s; 51 ˚C, 30s; 72 ˚C, 30s), and interrogating SYBR Green fluorescence after the anneal step of each cycle as previously described (2). Raw fluorescence data were then analyzed using CFX Maestro™ software.

**Aptamer library sequencing and analysis**

DNA libraries recovered after each selection round described above and the original, naïve library (6643) were subjected to PCR with the same protocol described above but with unmodified primers (6708 and 6711). The previously determined optimum number of cycles was used for each library to obtain unmodified, duplex DNA. PCR product size and quality were assessed by sample analysis by electrophoresis through 10% polyacrylamide gels with post-staining using SYBR Gold (Invitrogen, S11494) at this point and just before sequencing. MinElute spin columns (Qiagen, 28204) were used to purify the PCR products. A Qubit HS Duplex DNA Quantification kit (Invitrogen, Q32851) and Qubit 3.0 Fluorometer (Invitrogen, Q33216) were used to quantify the concentration of the purified PCR product. For each library, 10 ng was used as input into the NEBNext Ultra II DNA Library Prep with Sample Purification Beads (NEB, E7103S). Volumes recommended by the manufacturer were halved, and in the purification of adaptor-ligated DNA (step 3), Qiagen MinElute columns were used in place of beads. Libraries were barcoded using NEBNext Multiplex Oligos for Illumina (Index Primers Set 1) (NEB, E7335S). Paired end sequencing was performed for 100 cycles on a single lane of an Illumina MiSeq instrument with 30% PhiX DNA spike.

Usearch was used to merge paired end reads. Reads with a total quality score >0.5 were discarded (3). SeqKit was used to create uniform forward reads (4). AptaSUITE was used to filter any reads that did not contain both the forward and reverse primers within an error of 3 bases. AptaSUITE was also used to rank aptamers by their abundance and cluster the aptamers by sequence similarity (AptaCluster) (5).

**qPCR binding assay**

For 10-cm dish scale, 50 pmol library was prepared in 2 mL of wash buffer (50 nM). The solution was heated at ~95 ˚C for 5 min and snap cooled on ice for ten min before returning to room temperature. tRNA (100 µg/mL) and BSA (100 µg/mL) were then added as non-specific competitors. Media was aspirated from each dish of senescent MAFs and each was rinsed with warm DPBS once. Each library solution was added to a dish of senescent MAFs and incubated at 37 ˚C for 30 min. The senescent MAFs were washed on each dish 3× with 2 mL wash buffer. The cells were scraped into wash buffer and collected by centrifugation (500×g for 5 min). Supernatants were removed and cell pellets resuspended in 500 µL wash buffer each. Cells were then lysed by heating at 95 ˚C for 10 min. Debris was removed by centrifugation at 13,100×g for 5 min, and the supernatants containing the recovered libraries were moved to fresh tubes.

For 24-well plate scale, replicate wells were prepared for both senescent and control MAF conditions. Solutions were prepared as for the 10-cm dish scale above but with the candidate sequences rather than libraries. Media was aspirated from each well and each was rinsed with warm DPBS once. Candidate solutions were added to the wells (300 µL each well) and incubated at 37 ˚C for 40 min. Each well was washed 3× with 500 µL wash buffer. The cells were scraped into 300 µL wash buffer. Cells were then lysed by heating at 95 ˚C for 10 min.

qPCR reactions (30 µL) were performed with the following reagent volumes: 15 µL PerfeCTa SYBR Green FastMix for IQ (Quantabio, 95071-012), 3 µL 5 µM forward primer, 3 µL 5 µM reverse primer, 7.5 µL water, 1.5 µL template. Samples were transferred to a CFX96 Touch Deep Well Real-Time PCR Detection System (Bio-Rad Laboratories, 1854095) and subjected to thermal cycling using a protocol of 95 ˚C, 30s; 40×(95 ˚C 15s, 51 ˚C 30s, 72 ˚C 30s, 48 ˚C 30s), and interrogating SYBR Green fluorescence after the anneal step of each cycle as previously described (2). Raw fluorescence data were then analyzed using CFX Maestro™ software.

**Immunofluorescence staining and IncuCyte SX5 imaging**

Cells were washed with wash buffer (DPBS (Gibco) supplemented with 5mM $MgCl_2$ and 4.5 g/L glucose). Aptamer solution (wash buffer and 50 nM aptamer) was heated at 95 ˚C for 5 min and snap cooled on ice. Final aptamer solution (wash buffer supplemented with 50 nM aptamer, 10 mg/mL BSA, and 10 mg/mL tRNA) was added to cells for 45 min at 37°C. Cells were washed 3× with wash buffer and incubated with 100% methanol for 15 min at -20°C. Cells were washed 3× with wash buffer and incubated with Streptavidin, Alexa Fluor 647 conjugate (Invitrogen, S21374) secondary stain at 1:500 dilution in DPBS for 1 h at room temperature. Cells were washed 3× with DPBS (Gibco) and imaged with IncuCyte SX5

(Sartorius) at 10× magnification. Quantification of aptamer staining was calculated as total NIR signal intensity per image field, with nine replicates per condition.

**Proteomics**

Based on a previously described method, we used a SILAC-based mass spectrometry approach to identify molecular targets of two aptamers (6). C2C12 cells were cultured in media supplemented with heavy or light amino acids (Thermo Scientific, A33972) for five passages to ensure amino acid incorporation. These C2C12 cells were then plated in replicates and treated as described above for etoposide-induced senescence. Aptamers 6756 and 6762, and non-specific control oligonucleotide 6766 were each prepared at 100 nM in wash buffer, heated at 90 °C for 10 min, and snap cooled on ice. tRNA (100 µg/mL) and BSA (100 µg/mL) were then added as non-specific competitors. Medium was aspirated from senescent C2C12 cells, and the dish was rinsed with warm wash buffer once. The aptamer or negative control solutions were added to the senescent C2C12 cells and incubated at 37 °C for 40 min. Cells were washed twice with wash buffer and fixed with 2% formaldehyde at room temperature for 15 min. Cells were washed twice with PBS. The cells were scraped into lysis buffer (10 mM Tris-HCl pH 7.5, 10 mM NaCl, 0.5% IGEPAL) and incubated on ice for 1 h. Debris was removed by centrifugation at 8000 RCF for 5 min. The lysate supernatant was incubated with 50 µL M-270 Dynabeads (Invitrogen, 65305), that had been prepared according to manufacturer's instructions, for 2 h with end-over-end mixing at 4 °C. The beads were washed twice with lysis buffer, twice with PBS, and twice with water using a magnetic stand. The beads were resuspended in 50 µL elution buffer (10 mM Tris-HC pH 8, 10 mM

EDTA, 5 mM DTT, 1% SDS). Light and heavy samples were combined in duplicate as follows: light 6756 and heavy 6766, light 6762 and heavy 6766, light 6766 and heavy 6756, light 6766 and heavy 6762. The samples were heated at 95 ˚C for 1 h and a magnetic stand was used to remove the beads. The eluate was frozen at -80 ˚C until preparation for mass spectrometry analysis.

The samples were digested with Micro S-Traps (Protifi). The peptide extracts were analyzed by nano-ESI-LC/MS/MS with an Exploris mass spectrometer coupled to a Dionex nano-LC system (Thermo Scientific). The LC system used a gradient with solvent A (2% ACN, 0.2% formic acid, in water) and solvent B (80% ACN, 10% IPA, 0.2% formic acid, in water) as follows: -4-5 min, 5% B; 5- 125 min 5-45 % gradient; 125-128 min 45-95% gradient; 128-132 min 95% B; 132-134 min 95-5% B gradient; 134-137 min 5% B with a flow rate of 300 nL/min. The mass spectrometer had a resolution of 60,000 at 200 m/z and used data dependent acquisition, with a full MS1 scan ranging from 340-1600 m/z. Dynamic exclusion was set to 25 s. Cycle time was 3. All MS/MS spectra were analyzed using MaxQuant (Max Planck Institute of Biochemistry, Version 1.6.17.0) (7). Each was set up to search the current UniProt database, assuming trypsin digestion with up to two miscleavages with a fragment ion tolerance of 20 PPM and parent ion tolerance of 4.5 ppm (UniProt is provided in the public domain by the Swiss Institute of Bioinformatics, Geneva, Switzerland, www.uniprot.org). Oxidation of methionine was set as a variable modification, and carbamidomethylation of cysteine (iodoacetamide derivative) was set as a fixed modification, along with an allowance for the Arg +10 and Lys +8 Silac labels.

**RNA-seq**

Total RNA concentration and quality were determined using Qubit fluorometry (Invitrogen) and the Agilent BioAnalyzer using the Total RNA Pico chip (Agilent). cDNA libraries were prepared using 200 ng of total RNA according to the manufacturer's instructions for the TruSeq Stranded mRNA Sample Prep Kit (Illumina). The concentration and size distribution of the completed libraries were determined using an Agilent TapeStation D1000 and Qubit fluorometry. Libraries were sequenced at six samples per lane following the standard protocol for the Illumina NovaSeq™ 6000 and using the NovaSeq XP 2-Lane kit for individual lane loading. The flow cell was sequenced as 100 × 2 paired end reads using the NovaSeq SP sequencing kit and NovaSeq Control Software v1.7.5. Base-calling was performed using Illumina's RTA version 3.4.4.

RNA-Seq data were analyzed using the MAPRSeq pipeline v3.1.4 (8). Paired-end sequencing reads were mapped to the mouse genome (mm10) using STAR aligner (9). Gene and exon expression quantification were performed using the Subread package to obtain both raw and normalized (RPKM – Reads Per Kilobase per Million mapped reads) reads (10). Following alignment, secondary analyses, and quality control, the edgeR package v3.36.0 was used for obtaining differentially expressed genes (DEGs) (11). Genes were considered differentially expressed at a statistical significance threshold of p-value $< 0.05$ and an absolute $\log_2$ fold change value of greater than 1.

**Western blot**

Samples were prepared with NuPAGE LDS Sample Buffer (Invitrogen, NP0007) and NuPAGE Sample Reducing Agent (Invitrogen, NP0004). Heated samples at 95˚C for 3 min. Samples were subjected to electrophoresis at 150V for 1h on 10% Bis-Tris gel (Invitrogen, NO0303) with MES running buffer (Invitrogen, NP0002) and NuPAGE antioxidant (Invitrogen, NP0005). The gel was blotted onto a PVDF membrane (Bio-Rad, 1620174) using an XCell II Blot Module (Invitrogen, EI9051) in 1× NuPAGE transfer buffer (Invitrogen, NP0006) containing 20% methanol. Blotted membrane was briefly washed in TBST (50 mM Tris-HCl, pH 7.4, 150 mM NaCl, 0.1% Tween-20) and treated with blocking buffer (5% dry milk and 1 % BSA in TBST) overnight at 4˚C with gentle rocking. The membrane was incubated with an anti-fibronectin antibody (Novus Biologicals, NBP1-91258) diluted 1:1000 in blocking buffer overnight at 4˚C with gentle rocking. The membrane was washed with TBST four times for 15 min each at room temperature. The membrane was then incubated with an anti-rabbit secondary antibody with IRDye 680LT label (Licor Bio, 926-68021) diluted 1:10,000 in blocking buffer at room temperature for 1 h. The membrane was washed with TBST four times for 15 min each at room temperature. Membrane was analyzed with an Amersham Typhoon laser-scanner platform (Cytiva) using the IRshort channel.

**Protein capture assay**

Streptavidin Dynabeads M-270 (Invitrogen, 65305) were prepared with 50 µL beads (10 µg/mL) per aptamer and control oligonucleotide. Beads were mixed with 250 µL 2× B&W buffer. Tubes were placed on magnetic stand and supernatant was discarded. Beads were washed twice with 2× B&W buffer. Beads were loaded with aptamer or control oligonucleotides by adding 200 µL 2× B&W buffer, 50 µL water, and 200 pmol aptamer or control oligonucleotide followed by incubation at room temperature with end-over-end mixing for 1 h. Beads were washed three times with PBS supplemented with 5 mM $MgCl_2$ and 0.1% Tween-20. Native mouse fibronectin protein (Abcam, ab92784, 200 pmol) was added to each tube in PBS supplemented with 5 mM $MgCl_2$ and 0.1% Tween-20 and incubated with end-over-end mixing at room temperature for 40 min. Beads were washed three times with PBS supplemented with 5 mM $MgCl_2$ and 0.1% Tween-20. Bound protein was eluted by adding 13 µL PBS supplemented with 5 mM $MgCl_2$ and 0.1% Tween-20, 5 µL NuPAGE 4× LDS Sample Buffer (Invitrogen, NP0007), and 2 µL NuPAGE Sample Reducing Agent (Invitrogen, NP0004) followed by heating at 90 ˚C for 10 min and removal of beads using magnetic stand. Samples were analyzed by electrophoresis on 10% Bis-Tris gel (Invitrogen, NP0303) in MES running buffer (Invitrogen, NP0002) with NuPAGE antioxidant (Invitrogen, NP0005) followed by staining with Imperial Protein Stain (Thermo Scientific, 24615) and imaging with an Amersham Typhoon laser-scanner platform (Cytiva).

**Biolayer interferometry**

Biolayer interferometry was performed using an Octet Red96 (ForteBio) instrument. Octet Streptavidin biosensors (Sartorius, 18-5019) were used to immobilize biotinylated aptamers (500 nM). Native mouse fibronectin protein (Abcam, ab92784) was used as the analyte. Aptamers and fibronectin were diluted in 1× Octet Kinetics Buffer (Sartorius, 18-1105). The kinetic experiments were done at 30 ˚C. The experiment was performed with the following step parameters: baseline 180 s, loading 120 s, baseline 180 s, association 600 s, dissociation 600 s. Background subtracted sensograms were used to perform curve fitting with a 1:1 binding model.

**Cell culture confocal microscopy**

Control and senescent cells were prepared as described in the cell culture section on glass bottom dishes (MatTek, P356-1.5-14-C). Cells were washed once with wash buffer before staining with 50 nM aptamer or negative control oligonucleotide in binding buffer. The cells were then washed three times with PBS and fixed with formaldehyde at room temperature for 15 min. Then the cells were washed with PBS once and incubated with blocking buffer (PBS including 2% FBS) for 30 min at room temperature. After an additional wash with PBS, the cells were incubated with anti-fibronectin primary antibody (Novus Biologicals, NBP1-91258) diluted 1:400 at room temperature for 1 h. The cells were washed three times with PBS then incubated with both an anti-rabbit IgG with Alexa Fluor 488 label (Invitrogen, A-11011) diluted 1:400 and Streptavidin, Alexa Fluor 647 conjugate (Invitrogen, S21374) diluted 1:250 at room temperature for 1 h in blocking buffer. The cells were rinsed once with PBS and stained with DAPI at room temperature for 10 min. Following three additional washes with PBS, the cells were imaged with a Zeiss 780 LSM microscope.

Fibronectin staining was quantified in CellProfiler by measuring red channel intensity of representative fields with the MeasureImageIntensity function. Stain intensity was normalized to cell number by counting DAPI objects in each field.

**Tissue confocal microscopy**

Lung tissue collection and immunofluorescence staining were done as previously described with minor changes (12). Briefly, 10 μm thick lung sections were made from OCT embedded tissue and mounted on microscope slides. Sections were not permeabilized but were blocked with 5% BSA in PBS for 30 min at room temperature. Incubation with primary anti-fibronectin antibody (Abcam, ab2413) diluted 1:250 and 100 nM aptamer or negative control oligonucleotide was done overnight at 4˚C in binding buffer. The sections were washed twice with PBS followed by a 2-h incubation with the secondary antibody (Invitrogen, A11012) diluted 1:250 in PBS at room temperature. After two PBS washes, samples were stained with DAPI diluted 1:10,000 in PBS for 20 min at room temperature followed by an additional two washes in PBS. While in aqueous mounting media, the sections were sealed with a coverslip. All sections were stored at 4˚C until imaged.

A "prevention" strategy was designed to remove p16-positive senescent cells in INK-ATTAC mice as they arise with aging to limit their accumulation. Mice were treated with AP or vehicle starting at midlife (12 months) twice a week continuously until they reached 21 months of age. After 48 h past the last treatment dose, the lung tissues were collected to prepare for staining and imaging.

Images were collected by confocal microscopy using a Zeiss LSM 980. When entire tissue sections are shown, confocal images were taken at 5× with Z-stacks summed and tiled. Number of tiles were selected so that the tissue in its entirety was imaged, thus differed across slices. All 20× and 40× images were taken with Z-stacks summed, and the number of slices and slice size matching for a given experiment. Orthogonal projection was done in the frontal plane (xy).

Fibronectin and 6762 staining were quantified in CellProfiler by measuring red or green channel intensity of representative fields respectively with the MeasureImageIntensity function. Stain intensity was normalized to cell number by counting DAPI objects in each field.

**Table S1. Names and primary structures of primers, libraries, and aptamers.**

| Serial number | Oligonucleotide sequence[1] |
|---|---|
| 6638 | /56-FAM/TGCGTATTGACACATGCGTG |
| 6643 | TGCGTATTGACACATGCGTG$N_{40}$CAGGCATAGGTATGGGCTTT |
| 6644 | $A_{20}$/iSp9//iSp9/AAAGCCCATACCTATGCCTG |
| 6708 | TGCGTATTGACACATGCGTG |
| 6711 | AAAGCCCATACCTATGCCTG |
| 6756 | /56-FAM/TGCGTATTGACACATGCGTGAAACGGACTTTTATTAGACAAATAATGCATGT CTCGAATTCAGGCATAGGTATGGGCTTT/3BioTEG/ |
| 6757 | /56-FAM/TGCGTATTGACACATGCGTGGCCCAAATGATATTGAGCAATATCATGGTAGG TAAGCTCGCAGGCATAGGTATGGGCTTT/3BioTEG/ |
| 6758 | /56-FAM/TGCGTATTGACACATGCGTGAAAACCCGGCGTTATTGCTGTAATATTAGCCT GGGAGTTTCAGGCATAGGTATGGGCTTT/3BioTEG/ |
| 6759 | /56-FAM/TGCGTATTGACACATGCGTGGAGAGTCGATCAATTAGCTTAGGTATTAATTC TTCGCGGACAGGCATAGGTATGGGCTTT/3BioTEG/ |
| 6760 | /56-FAM/TGCGTATTGACACATGCGTGCATAGGTATCCGCAATCCGTTATTCAGAGAAT ATTAGGATCAGGCATAGGTATGGGCTTT/3BioTEG/ |
| 6761 | /56-FAM/TGCGTATTGACACATGCGTGGTGTTATCATAATGCGGTATTGACCTTTATAA TATTAATCCAGGCATAGGTATGGGCTTT/3BioTEG/ |
| 6762 | /56-FAM/TGCGTATTGACACATGCGTGGCTTAATGCAATCCATCTGTTGCATTTTTCTA AACGTCGCCAGGCATAGGTATGGGCTTT/3BioTEG/ |
| 6763 | /56-FAM/TGCGTATTGACACATGCGTGCTATCGTTATGCACCGAATATTAGATAATGCA ATACGCCTCAGGCATAGGTATGGGCTTT/3BioTEG/ |
| 6764 | /56-FAM/TGCGTATTGACACATGCGTGTAGCAAAAGAGTCTCATATCCCGCTGGATATT ACGCGGTCCAGGCATAGGTATGGGCTTT/3BioTEG/ |
| 6765 | /56-FAM/TGCGTATTGACACATGCGTGATCGTGGACCATAGGTATCATTGTAAATACGG GCGTATTTCAGGCATAGGTATGGGCTTT/3BioTEG/ |
| 6766 | /56-FAM/TGCGTATTGACACATGCGTGCTTGTTATATAACGTGTCATAGCGTCCAAAAA GAAGTACACAGGCATAGGTATGGGCTTT/3BioTEG/ |
| 6767 | /56-FAM/TGCGTATTGACACATGCGTGTTGTACAATGGTCTATGATAATTATCAATAAG GTTTATGACAGGCATAGGTATGGGCTTT/3BioTEG/ |

[1]Modification abbreviations: /56-FAM/ is an isomer derivative of fluorescein with a six-carbon spacer at the 5' terminus of the oligonucleotide. /iSp9/ is an internal triethylene glycol spacer that creates a gap that is not extendable by Taq polymerase during PCR, allowing denaturing electrophoretic gel purification of the top DNA single strand PCR product. /3BioTEG/ is a biotin with an extended triethylene glycol spacer conjugated at the 3' oligonucleotide terminus.

**Table S2. Statistical comparison of aptamer binding in control vs senescent conditions.**

| Cell | Oligonucleotide | Method | p-value[1] | Significance[2] |
|---|---|---|---|---|
| MAF | 6756 | qPCR | <0.000001 | *** |
| | 6757 | qPCR | <0.000001 | *** |
| | 6758 | qPCR | <0.000001 | *** |
| | 6759 | qPCR | <0.000001 | *** |
| | 6760 | qPCR | <0.000001 | *** |
| | 6761 | qPCR | <0.000001 | *** |
| | 6762 | qPCR | <0.000001 | *** |
| | 6763 | qPCR | <0.000001 | *** |
| | 6764 | qPCR | 0.012706 | * |
| | 6765 | qPCR | 0.000005 | *** |
| | 6766 | qPCR | 0.737811 | n.s. |
| | 6767 | qPCR | 0.313484 | n.s. |
| MAF | 6756 | Imaging | <0.000001 | *** |
| | 6757 | Imaging | 0.066085 | n.s. |
| | 6758 | Imaging | <0.000001 | *** |
| | 6759 | Imaging | <0.000001 | *** |
| | 6760 | Imaging | <0.000001 | *** |
| | 6761 | Imaging | <0.000001 | *** |
| | 6762 | Imaging | <0.000001 | *** |
| | 6763 | Imaging | 0.000169 | *** |
| | 6764 | Imaging | 0.659772 | n.s. |
| | 6765 | Imaging | 0.035911 | * |
| | 6766 | Imaging | 0.766286 | n.s. |

| | | | | |
|---|---|---|---|---|
| | 6767 | Imaging | 0.807914 | n.s. |
| MAF serum starved | 6756 | Imaging | <0.000001 | *** |
| | 6757 | Imaging | 0.000195 | *** |
| | 6758 | Imaging | <0.000001 | *** |
| | 6759 | Imaging | <0.000001 | *** |
| | 6760 | Imaging | <0.000001 | *** |
| | 6761 | Imaging | <0.000001 | *** |
| | 6762 | Imaging | <0.000001 | *** |
| | 6763 | Imaging | <0.000001 | *** |
| | 6764 | Imaging | 0.390611 | n.s. |
| | 6765 | Imaging | 0.006236 | ** |
| | 6766 | Imaging | 0.192352 | n.s. |
| | 6767 | Imaging | 0.319937 | n.s. |
| MAF replated | 6756 | Imaging | 0.050334 | n.s. |
| | 6757 | Imaging | 0.547740 | n.s. |
| | 6758 | Imaging | 0.000103 | *** |
| | 6759 | Imaging | 0.023875 | * |
| | 6760 | Imaging | 0.097555 | n.s. |
| | 6761 | Imaging | <0.000001 | *** |
| | 6762 | Imaging | 0.025313 | * |
| | 6763 | Imaging | 0.126055 | n.s. |
| | 6764 | Imaging | 0.589578 | n.s. |
| | 6765 | Imaging | 0.093085 | n.s. |
| | 6766 | Imaging | 0.314707 | n.s. |
| | 6767 | Imaging | 0.379835 | n.s. |
| C2C12 | 6756 | Imaging | <0.000001 | *** |

| | | | |
|---|---|---|---|
| 6757 | Imaging | 0.337705 | n.s. |
| 6758 | Imaging | 0.000036 | *** |
| 6759 | Imaging | <0.000001 | *** |
| 6760 | Imaging | <0.000001 | *** |
| 6761 | Imaging | <0.000001 | *** |
| 6762 | Imaging | <0.000001 | *** |
| 6763 | Imaging | 0.005323 | ** |
| 6764 | Imaging | 0.395057 | n.s. |
| 6765 | Imaging | 0.118345 | n.s. |
| 6766 | Imaging | 0.165900 | n.s. |
| SA only | Imaging | 0.268197 | n.s. |

[1]Unpaired t-test with single pooled variance.

[2]*: p<0.05, **: p<0.01, ***: p<0.001

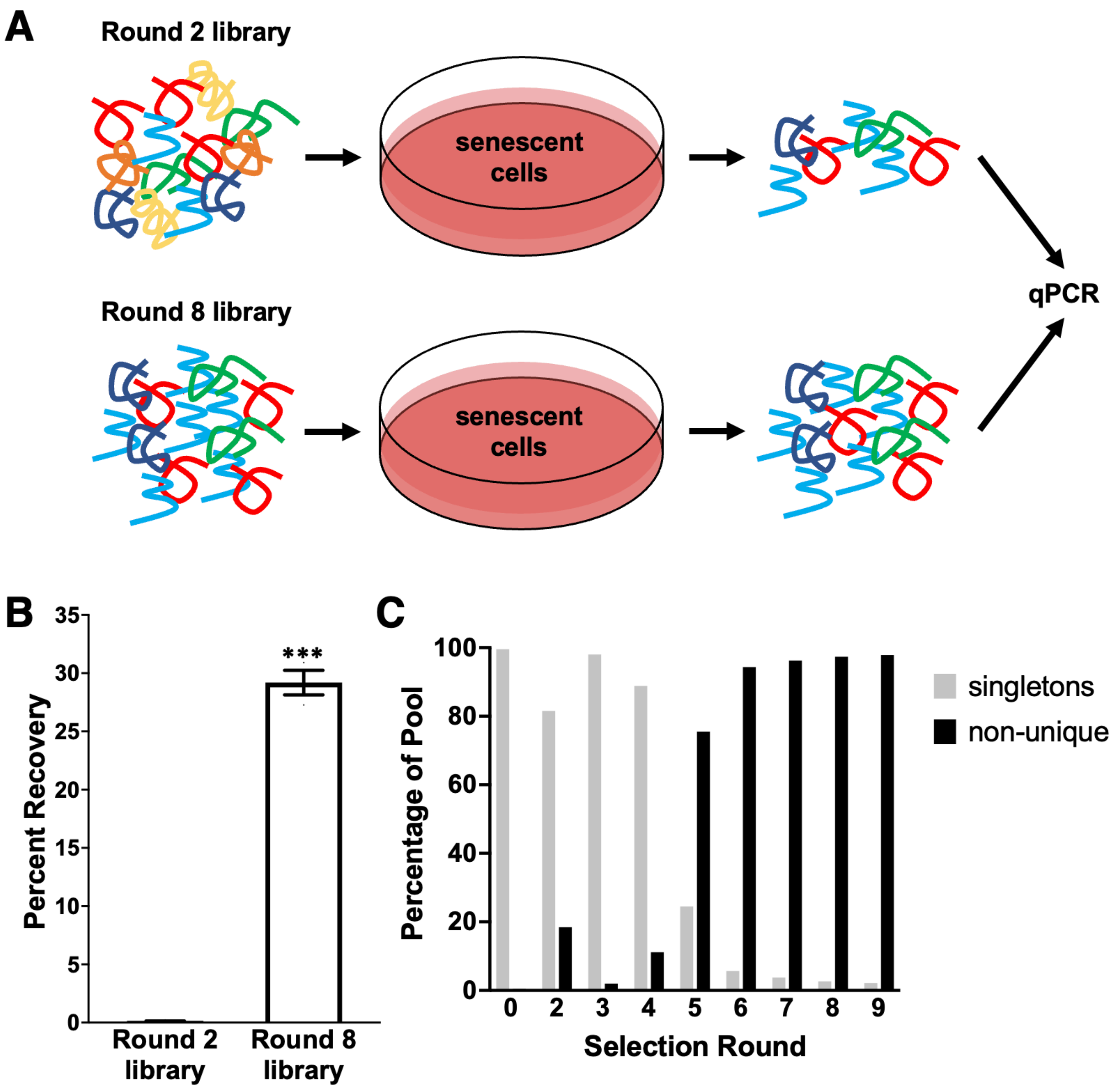

**Figure S1. Verification of overall pool behavior and deep sequencing data.** A) Schematic of experiment comparing binding of round 2 and round 8 libraries to senescent cells. B) qPCR quantification from comparison of round 2 and round 8 libraries (50 nM) binding to senescent cells. Error bars indicate standard error of technical replicates. C) Assessment of the overall enrichment of libraries in the deep sequencing data across rounds of selection. “Singletons” are sequences found only once in deep sequencing data, while “non-unique” refers to sequences found two or more times.

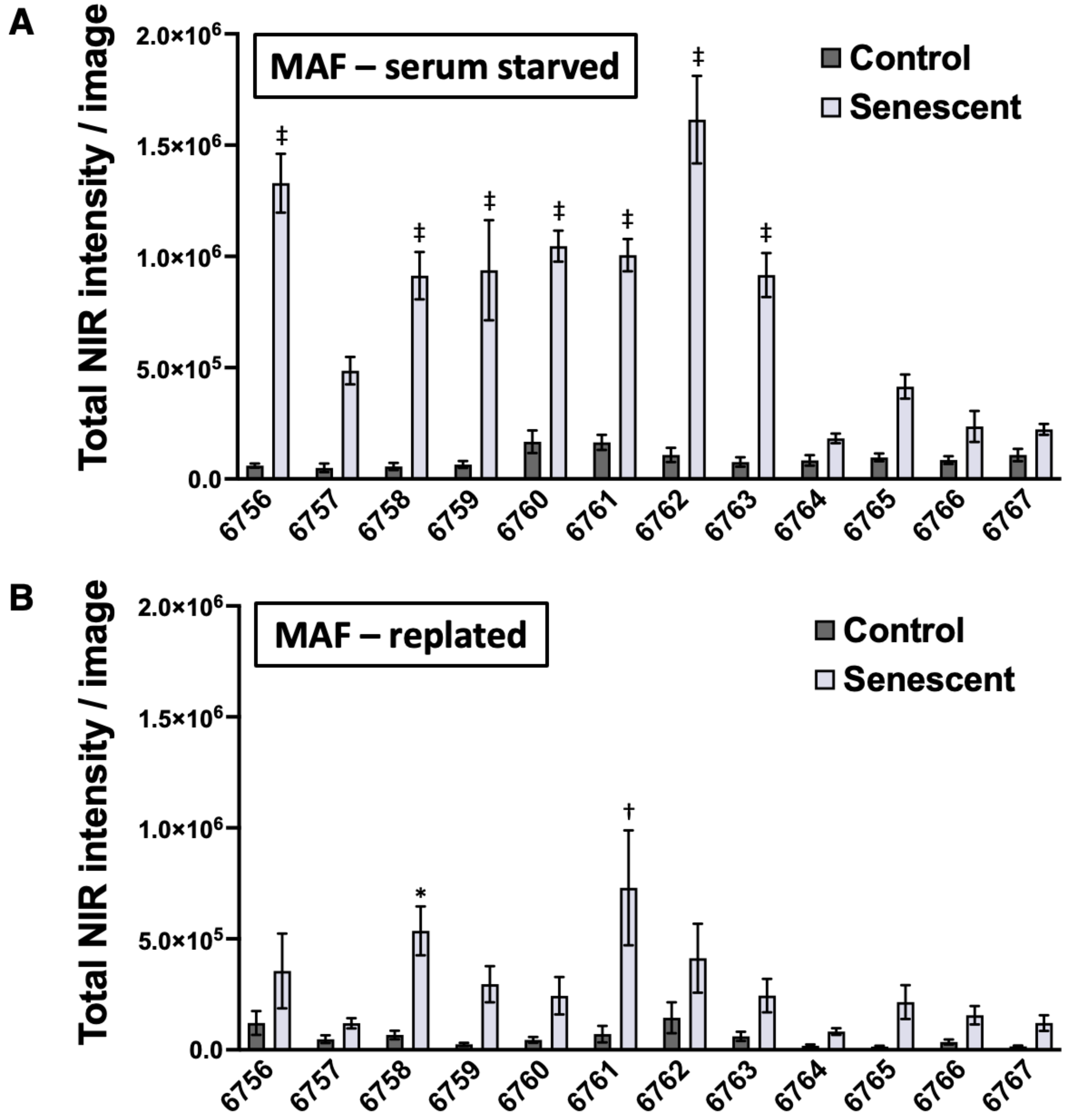

**Figure S2. Effects of cell culture time vs. replating on aptamer binding.** Quantification of 50 nM aptamer staining detected by secondary stain with AlexaFluor647 streptavidin is calculated as total NIR signal intensity per image field. Statistical significance is shown for candidates compared with control 6766 by one-way ANOVA with Dunnett's post-hoc test for multiple comparisons (*: $p<0.05$, †: $p<0.005$, ‡: $p<0.0005$). Error bars are shown as standard error for multiple image fields (n=9-16). A) Data for control MAFs that had been serum starved to match the plating time with senescent MAFs. B) Data for senescent MAFs that had been replated the day before staining to match the plating time of the control MAFs.

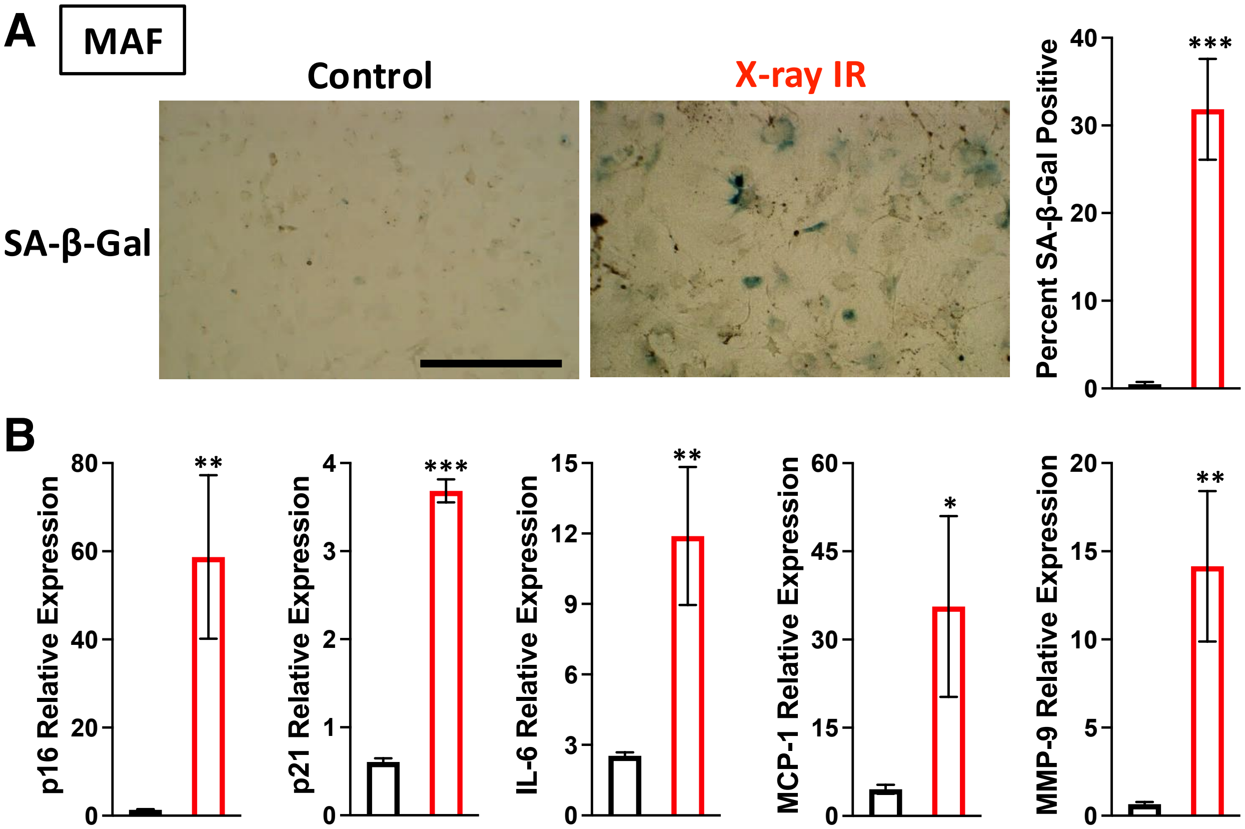

**Figure S3. Validation of senescent phenotype in X-ray irradiated MAFs.** A) Bright field images of control and X-ray irradiated MAFs show morphological changes and SA–β-Gal staining. The percent SA–β-Gal positive cells are quantified per field and compared using a two-tailed unpaired t-test (n=5; *p<0.05, **p<0.01, ***p<0.001). Scale bar is 400 μm. B) RT-qPCR of various markers of senescence assessed by ΔΔCT method normalized to TATA-box binding protein (TBP) as a reference gene. Each comparison is made using a two-tailed unpaired t-test (n=6; *p<0.05, **p<0.01, ***p<0.001).

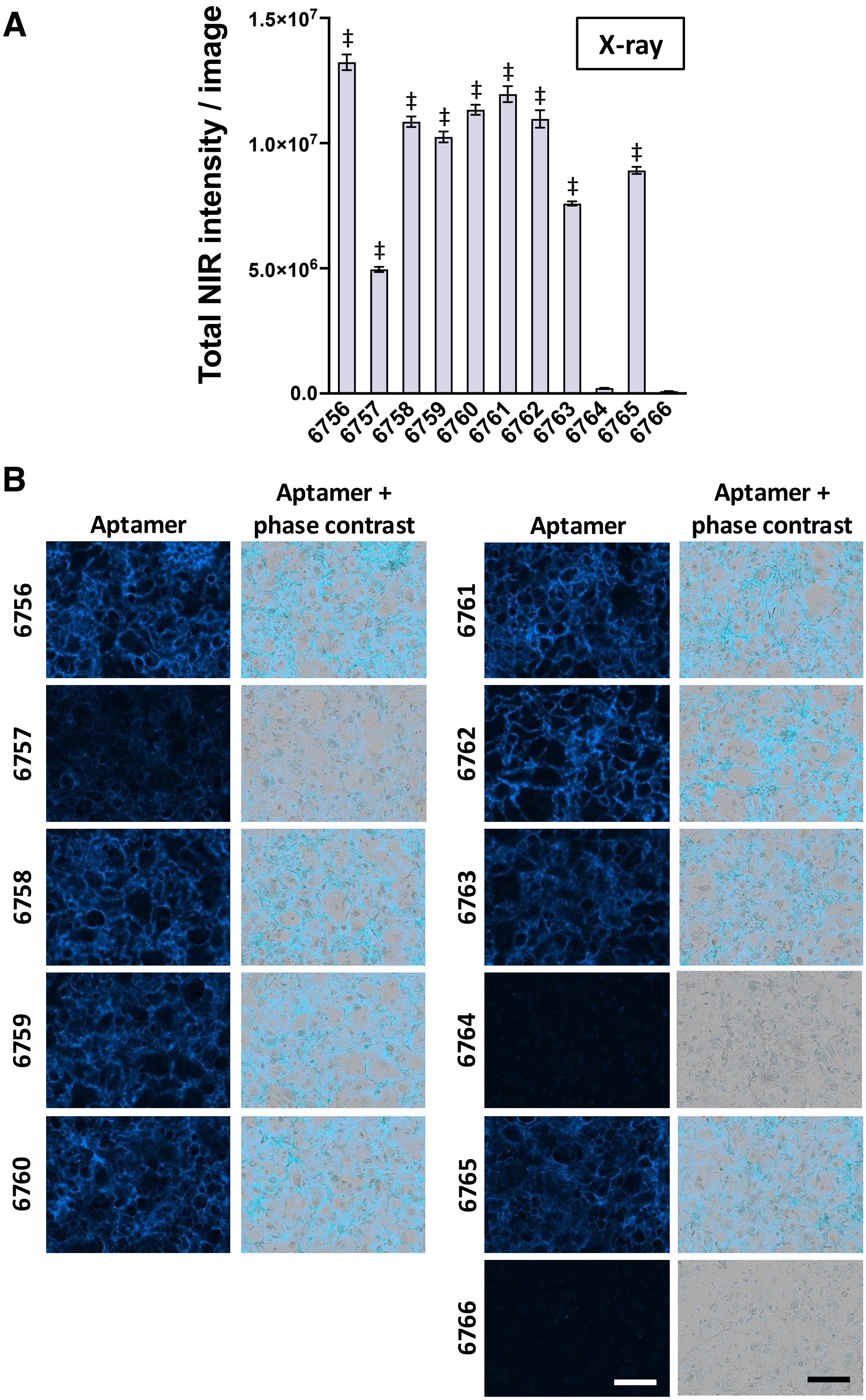
A
Total NIR intensity / image
1.5×10^7
1.0×10^7
5.0×10^6
0.0
X-ray
‡
B
Aptamer
Aptamer + phase contrast

**Figure S4. Screening candidate aptamer binding to X-ray-induced senescent MAFs.** A) Quantification of 50 nM aptamer staining was calculated as total NIR signal intensity per image field. Statistical significance is shown for candidates compared with control 6766 by one-way ANOVA with Dunnett's post-hoc test for multiple comparisons (*: p<0.05, †: p<0.005, ‡: p<0.0005). Error bars are shown as standard error for multiple image fields (n=16). B) Images taken with IncuCyte SX5 are shown at 10× magnification. Aptamer staining is shown in blue (secondary stain with AlexaFluor647 labeled streptavidin binding biotinylated candidates and control). Scale bars (lower right panels) are 400 μm.

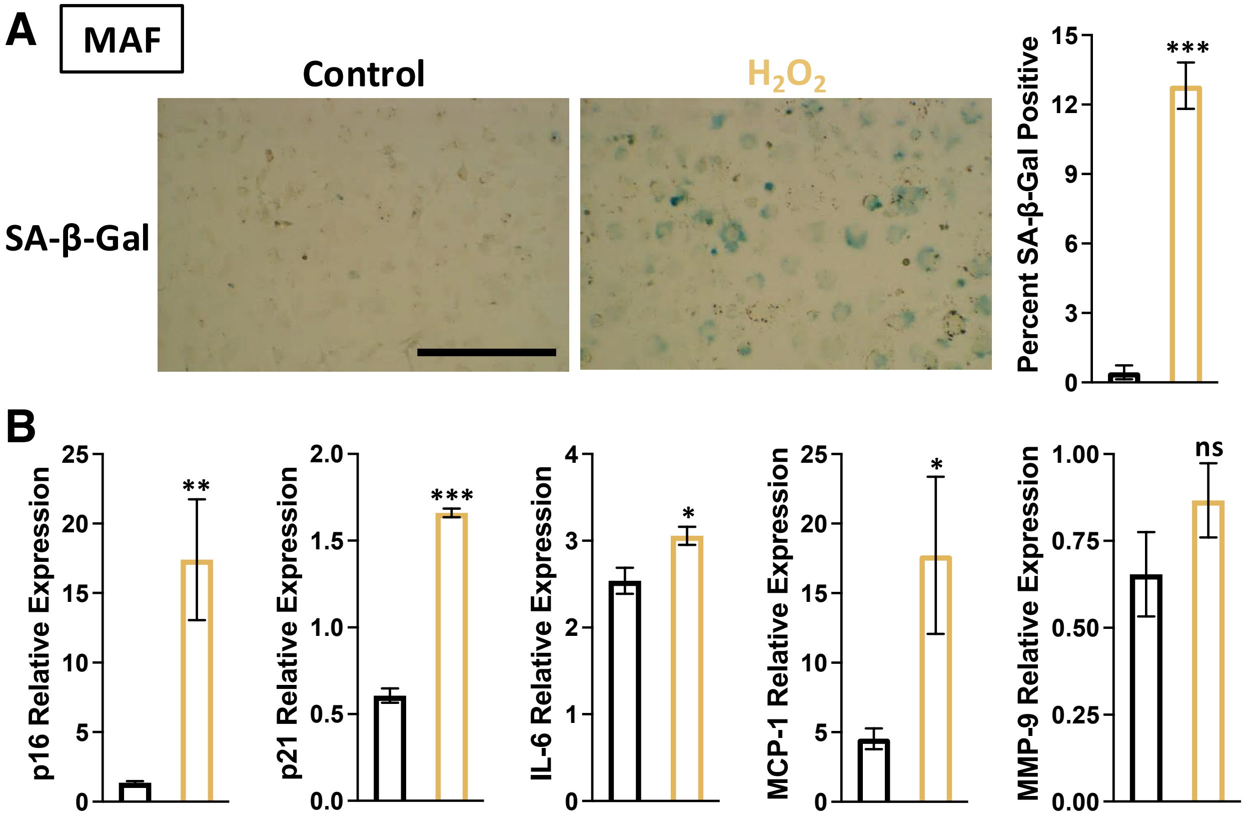

**Figure S5. Validation of senescent phenotype in hydrogen peroxide treated MAFs.** A) Bright field images of control and hydrogen peroxide treated MAFs show morphological changes and SA–β-Gal staining. The percent SA–β-Gal positive cells are quantified per field and compared using a two-tailed unpaired t-test (n=5; *p<0.05, **p<0.01, ***p<0.001). Scale bar is 400 μm. B) RT-qPCR of various markers of senescence assessed by ΔΔCT method normalized to TATA-box binding protein (TBP) as a reference gene. Each comparison is made using a two-tailed unpaired t-test (n=6; *p<0.05, **p<0.01, ***p<0.001).

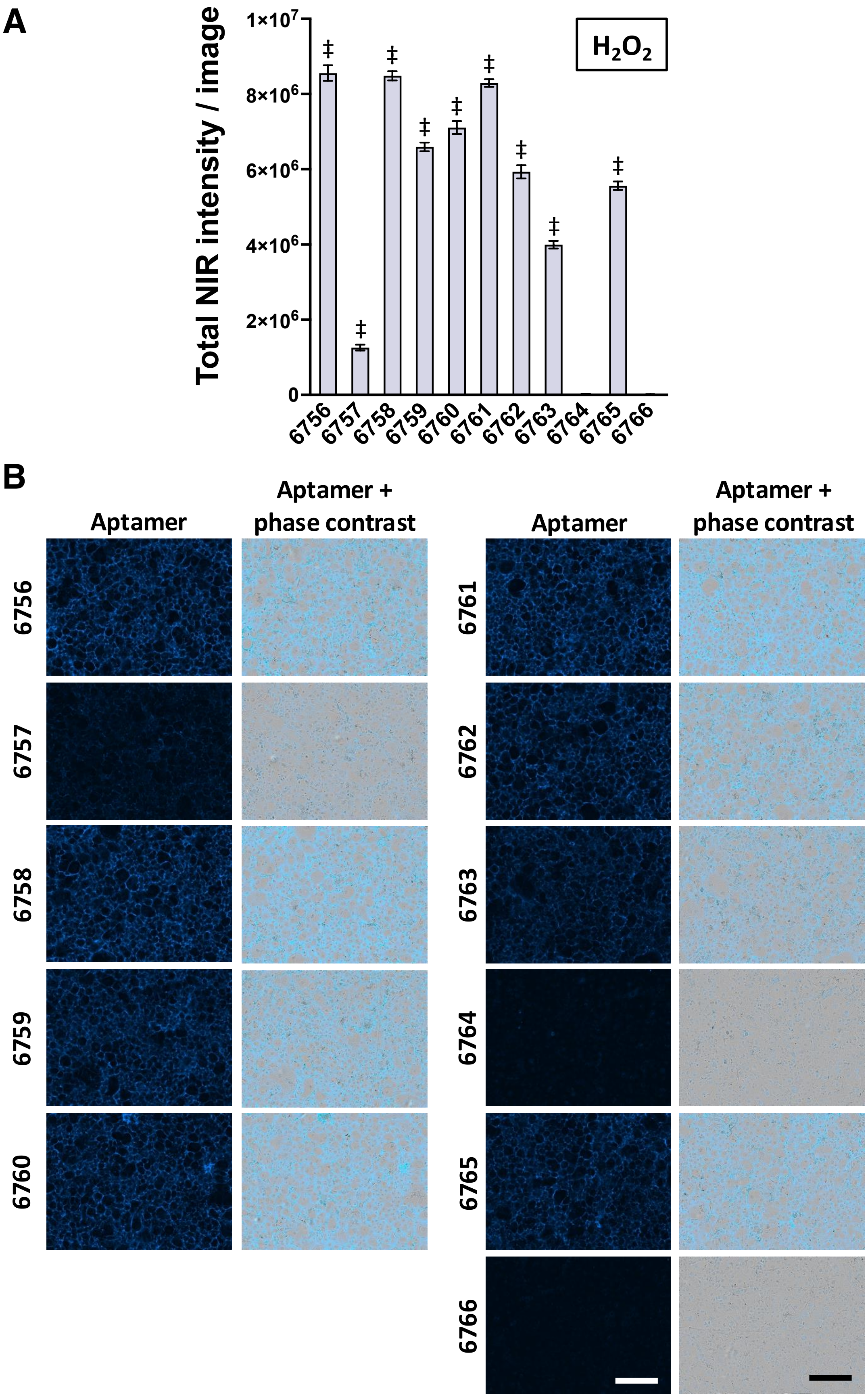
A
Total NIR intensity / image
1×10^7
8×10^6
6×10^6
4×10^6
2×10^6
0
H2O2
B
Aptamer
Aptamer + phase contrast
Aptamer
Aptamer + phase contrast

**Figure S6. Screening candidate aptamer binding to hydrogen peroxide-induced senescent MAFs.** A) Quantification of 50 nM aptamer staining was calculated as total NIR signal intensity per image field. Statistical significance is shown for candidates compared with control 6766 by one-way ANOVA with Dunnett's post-hoc test for multiple comparisons (*: $p<0.05$, †: $p<0.005$, ‡: $p<0.0005$). Error bars are shown as standard error for multiple image fields (n=16). B) Images taken with IncuCyte SX5 are shown at 10× magnification. Aptamer staining is shown in blue (secondary stain with AlexaFluor647 labeled streptavidin binding biotinylated candidates and control). Scale bars (lower right panels) are 400 μm.

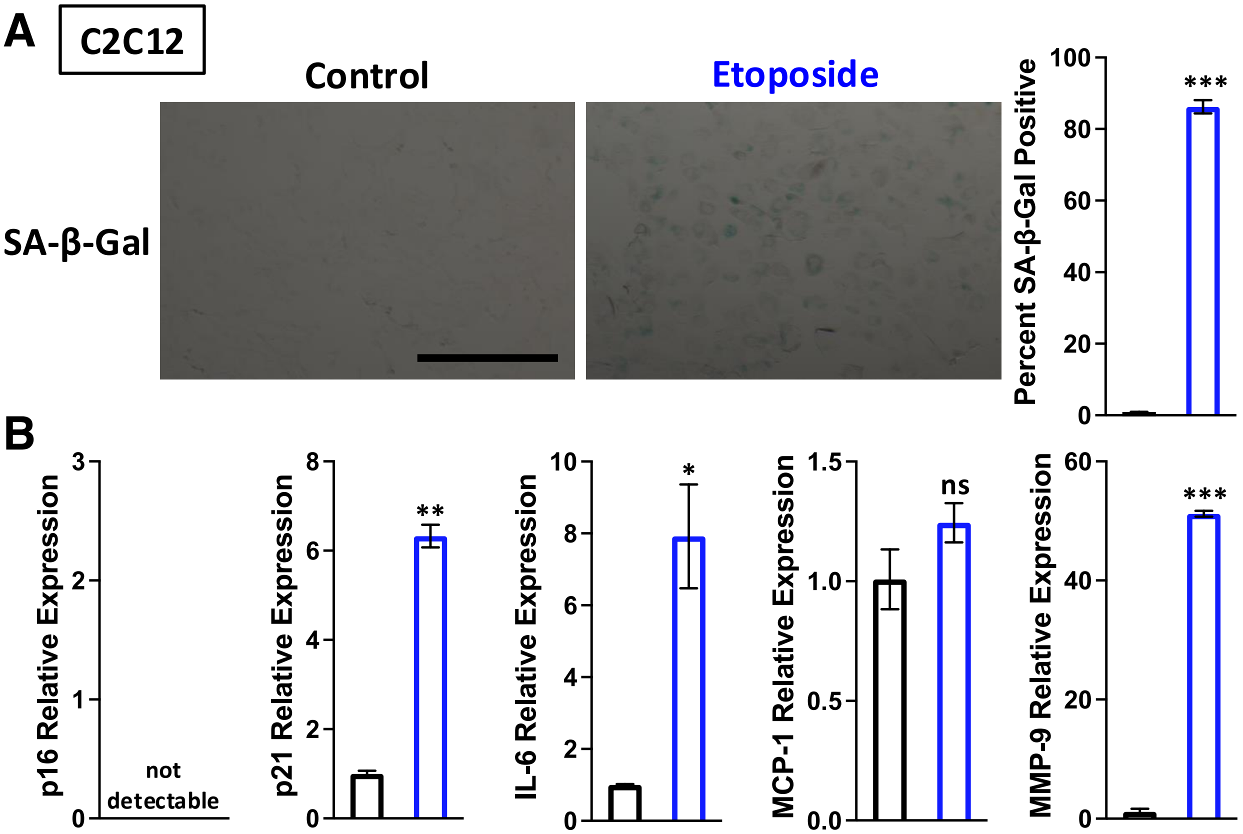

**Figure S7. Validation of senescent phenotype in etoposide-treated C2C12 cells.** A) Bright-field images of control and etoposide treated C2C12 cells show morphological changes and SA–β-Gal staining. The percent SA–β-Gal positive cells is quantified per field and compared with two-tailed unpaired t-test (n=6; *: $p<0.05$, **: $p<0.01$, ***: $p<0.001$). Scale bar is 400 μm. B) RT-qPCR of various markers of senescence assess by ΔΔCT method normalized to TATA-box binding protein (TBP) as a reference gene. Each comparison is made using a two-tailed unpaired t-test (n=3; *: $p<0.05$, **: $p<0.01$, ***: $p<0.001$).

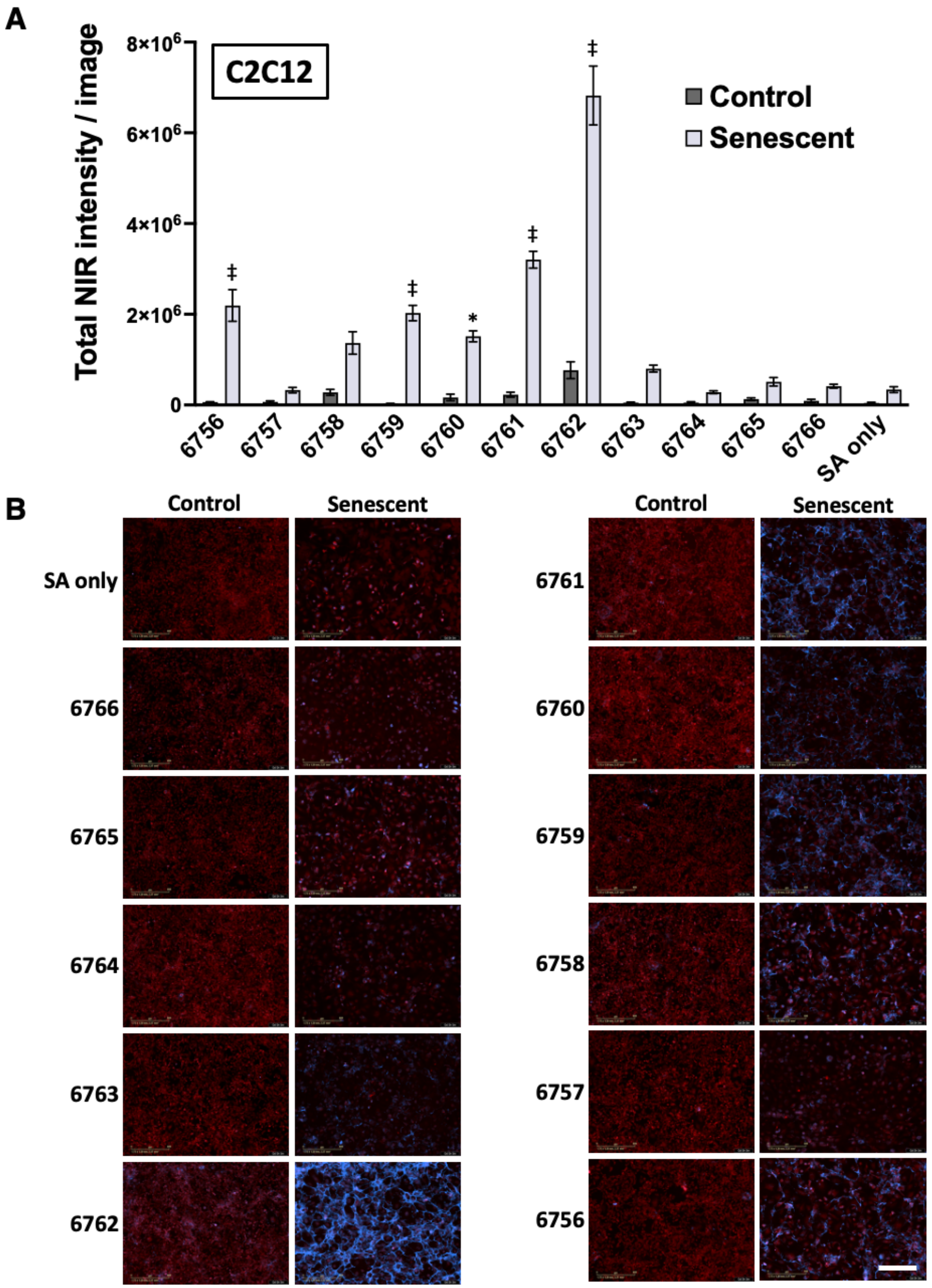
A
C2C12
Total NIR intensity / image
8×10^6
6×10^6
4×10^6
2×10^6
0
Control
Senescent
‡
‡
*
‡
‡
SA only
B
Control
Senescent
SA only
6766
6765
6764
6763
6762
Control
Senescent
6761
6760
6759
6758
6757
6756

**Figure S8. Screening candidate aptamer binding to senescent C2C12 cells.** A) Quantification of staining by 50 nM aptamer was calculated as total NIR signal intensity per image field. Statistical significance is shown for candidates compared with negative control 6766 by one-way ANOVA with Dunnett's post-hoc test for multiple comparisons (*: p<0.05, †: p<0.005, ‡: p<0.0005). Error bars are shown as standard error for multiple image fields (n=9). B) Images taken with IncuCyte SX5 are shown at 10× magnification. Constitutively expressed TdTomato is shown in red, and aptamer staining is shown in blue (secondary stain with AlexaFluor647 labeled streptavidin binding biotinylated candidates and controls). Scale bar (lower right panel) is 400 µm.

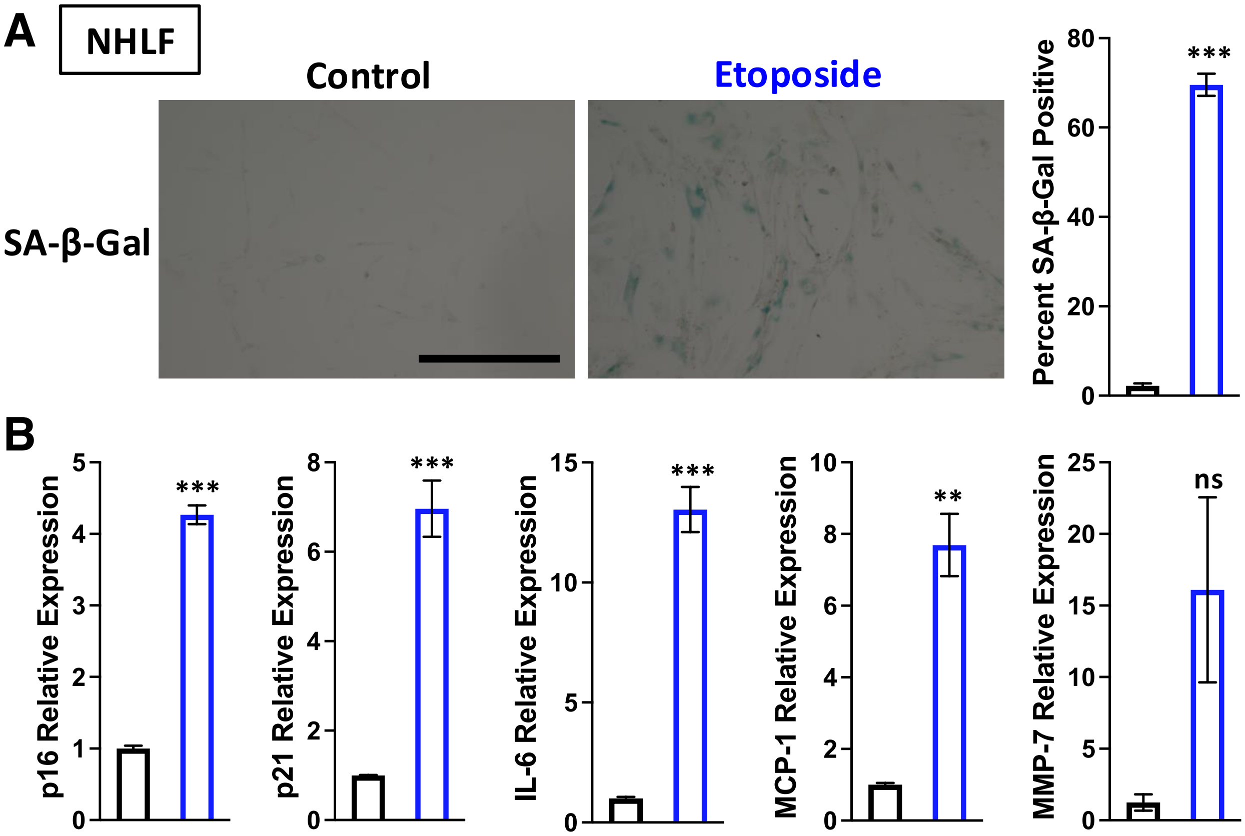

**Figure S9. Validation of senescent phenotype in etoposide-treated NHLF cells.** A) Bright field images of control and etoposide treated NHLF cells show morphological changes and SA–β-Gal staining. The percent SA–β-Gal positive cells is quantified per field and compared using a two-tailed unpaired t-test (n=10; *: p<0.05, **: p<0.01, ***: p<0.001). Scale bar is 400 µm. B) RT-qPCR of various markers of senescence assess by ΔΔCT method normalized to TATA-box binding protein (TBP) as a reference gene. Each comparison is made using a two-tailed unpaired t-test (n=3; *: p<0.05, **: p<0.01, ***: p<0.001).

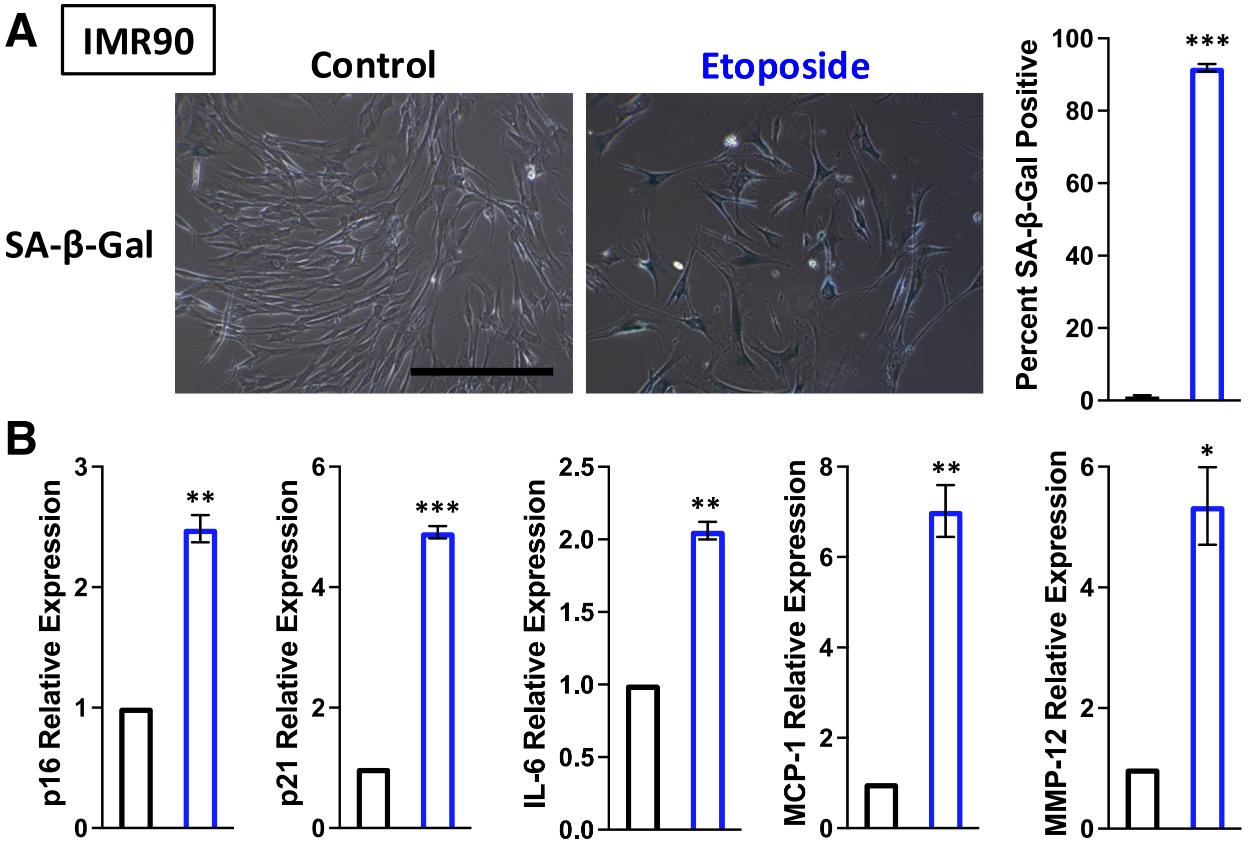

**Figure S10. Validation of senescent phenotype in etoposide-treated IMR90 cells.** A) Bright field images of control and etoposide treated IMR90 cells show morphological changes and SA–β-Gal staining. The percent SA–β-Gal positive cells is quantified per field and compared using a two-tailed unpaired t-test (n=5; *: p<0.05, **: p<0.01, ***: p<0.001). Scale bar is 400 μm. B) RT-qPCR of various markers of senescence assess by ΔΔCT method normalized to TATA-box binding protein (TBP) as a reference gene. Each comparison is made using a two-tailed unpaired t-test (n=3; *:p<0.05, **: p<0.01, ***: p<0.001).

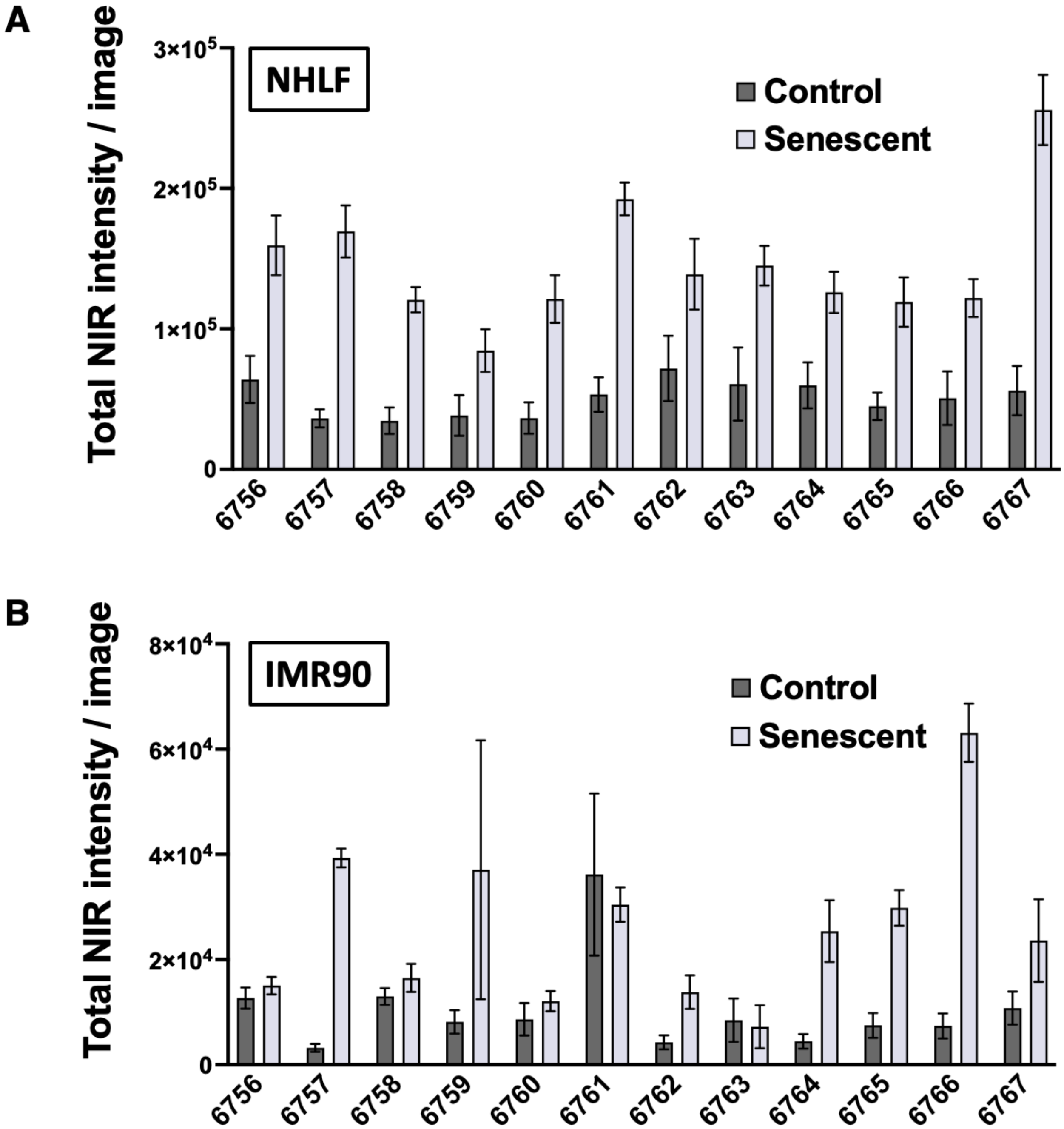

**Figure S11. Screening candidate aptamer binding to senescent NHLF and IMR90 cells.** A) Quantification of 50 nM aptamer staining of NHLF cells was calculated as total NIR signal intensity per image field. Error bars are shown as standard error for multiple image fields (n=9). B) Quantification of 50 nM aptamer staining of IMR90 cells was calculated as total NIR signal intensity per image field. Error bars are shown as standard error for multiple image fields (n=9).

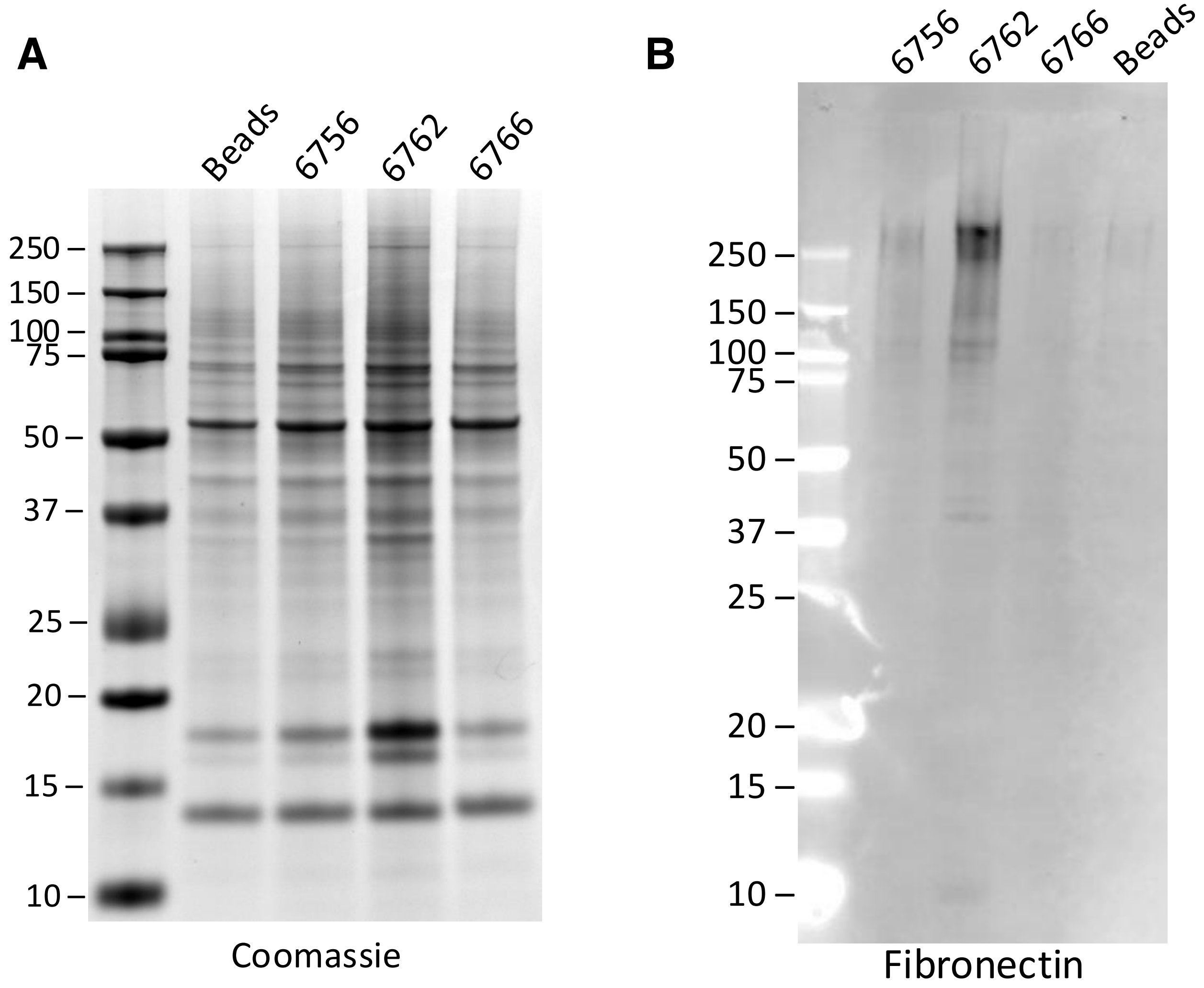

**Figure S12. Quality control data for anti-fibronectin Western blot.** A) Coomassie stained gel matched to show loading before transfer to membrane for Western blot indicates similar total loading which agrees with Qubit protein assay. B) The Western blot shown in Fig. 4A is shown uncropped here.

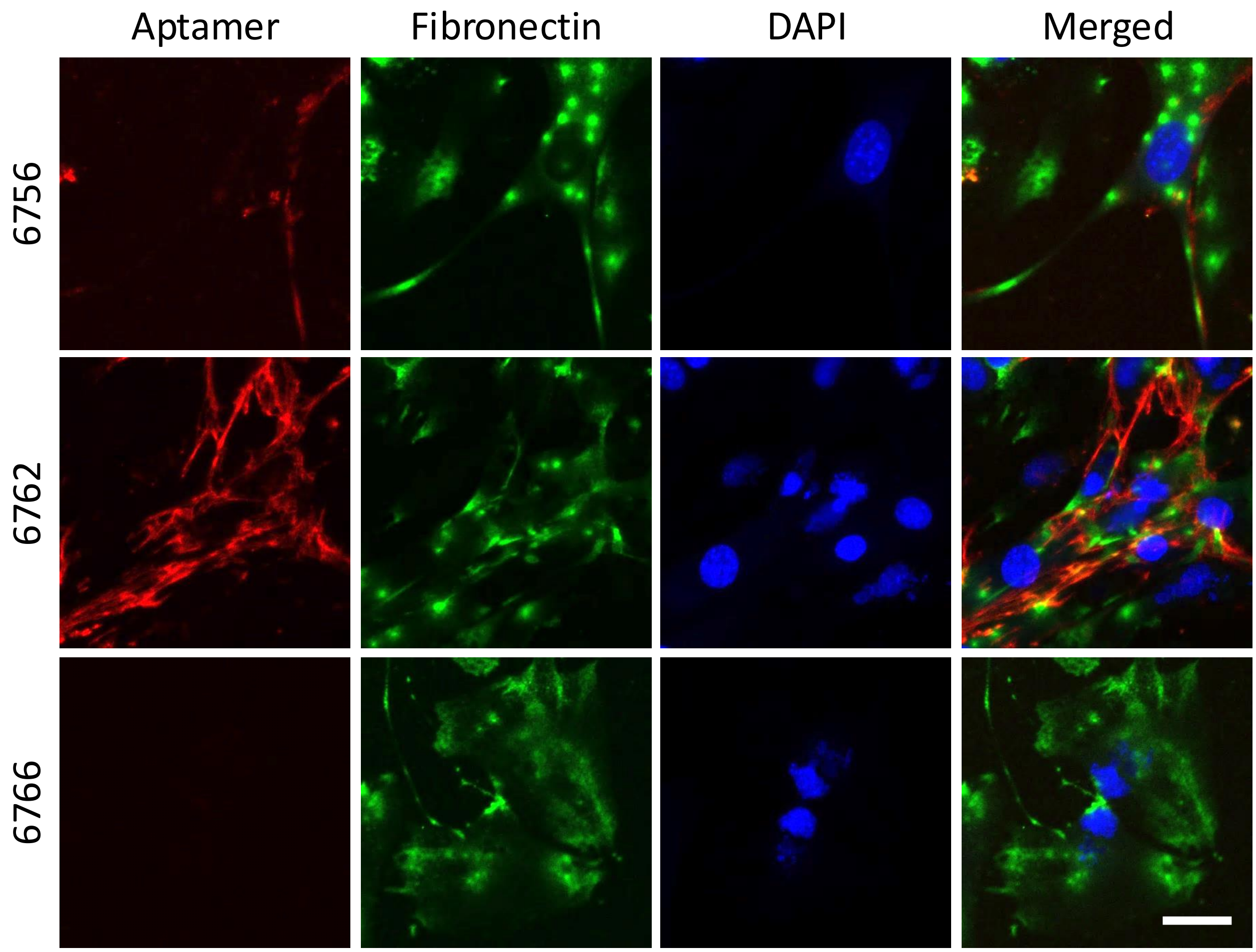

**Figure S13. Confocal imaging of senescent C2C12 cells co-stained with aptamers and anti-fibronectin antibody do not show complete overlap.** Biotinylated aptamers 6756 and 6762 and negative control 6766 are detected with secondary AlexaFluor647 labeled streptavidin (red). Primary anti-fibronectin antibody is detected by AlexaFluor568 labeled anti-rabbit secondary (green). Nuclei are stained with DAPI (blue). Scale bar (lower right panel) is 50 μm.

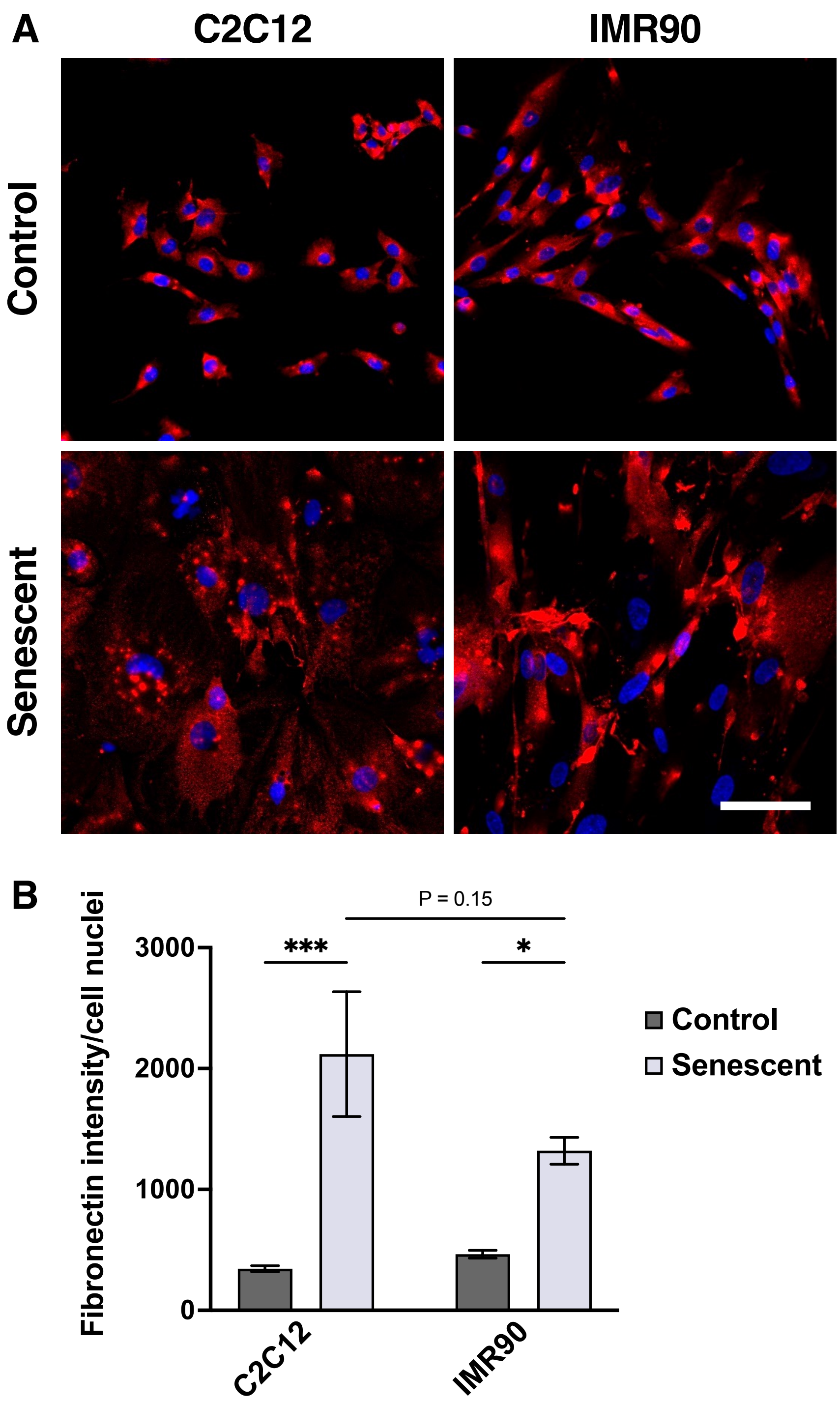

**Figure S14. Fibronectin staining is elevated in both senescent C2C12 and IMR90 cells compared with control cells.** A) Confocal images show anti-fibronectin antibody detected with AlexaFluor568 labeled anti-rabbit secondary (red). Nuclei are stained with DAPI (blue). B) Quantification of fibronectin staining intensity per cell nuclei taken from 9 image fields per condition. Each comparison is made using a two-tailed unpaired t-test (*$p<0.05$, **$p<0.01$, ***$p<0.001$). Scale bar (lower right panel) is 100 μm.

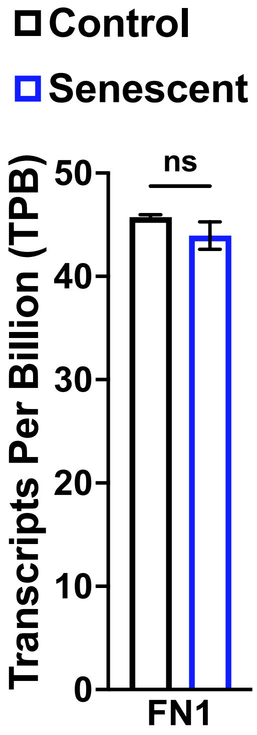

**Figure S15. RNA-seq relative quantitation of *FN1* transcripts.** Plot shows the mean transcripts per billion and standard error bars for mRNAs encoding *FN1* (n=3 each condition). Comparison is made using a two-tailed unpaired t-test (p = 0.25).

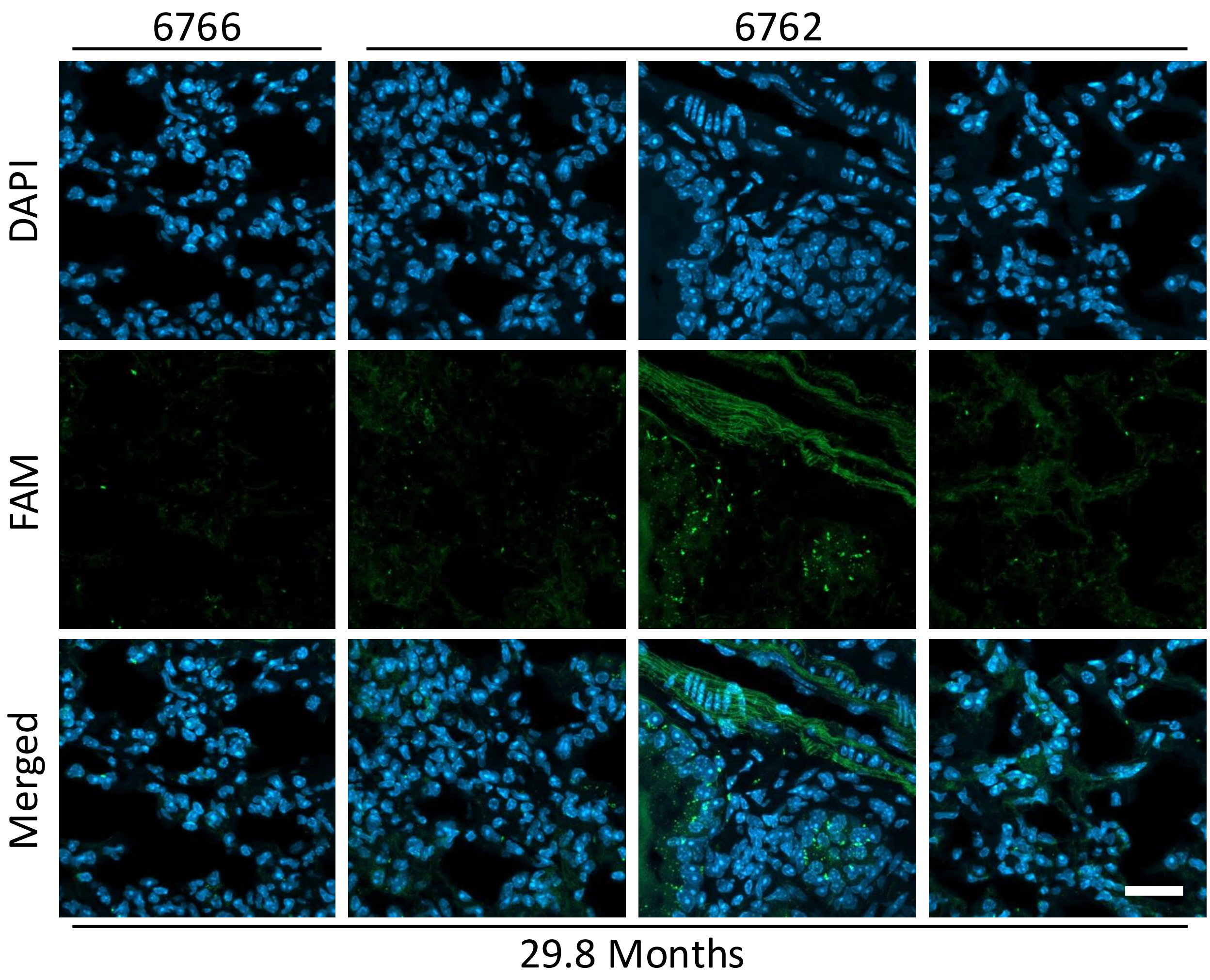

**Figure S16. Aptamer 6762 staining of naturally aged mouse lung tissue varies by region.** Aptamer 6762 and control 6766 are detected by fluorescein label (green). Nuclei are stained with DAPI (blue). Scale bar (lower right panel) is 25 μm.

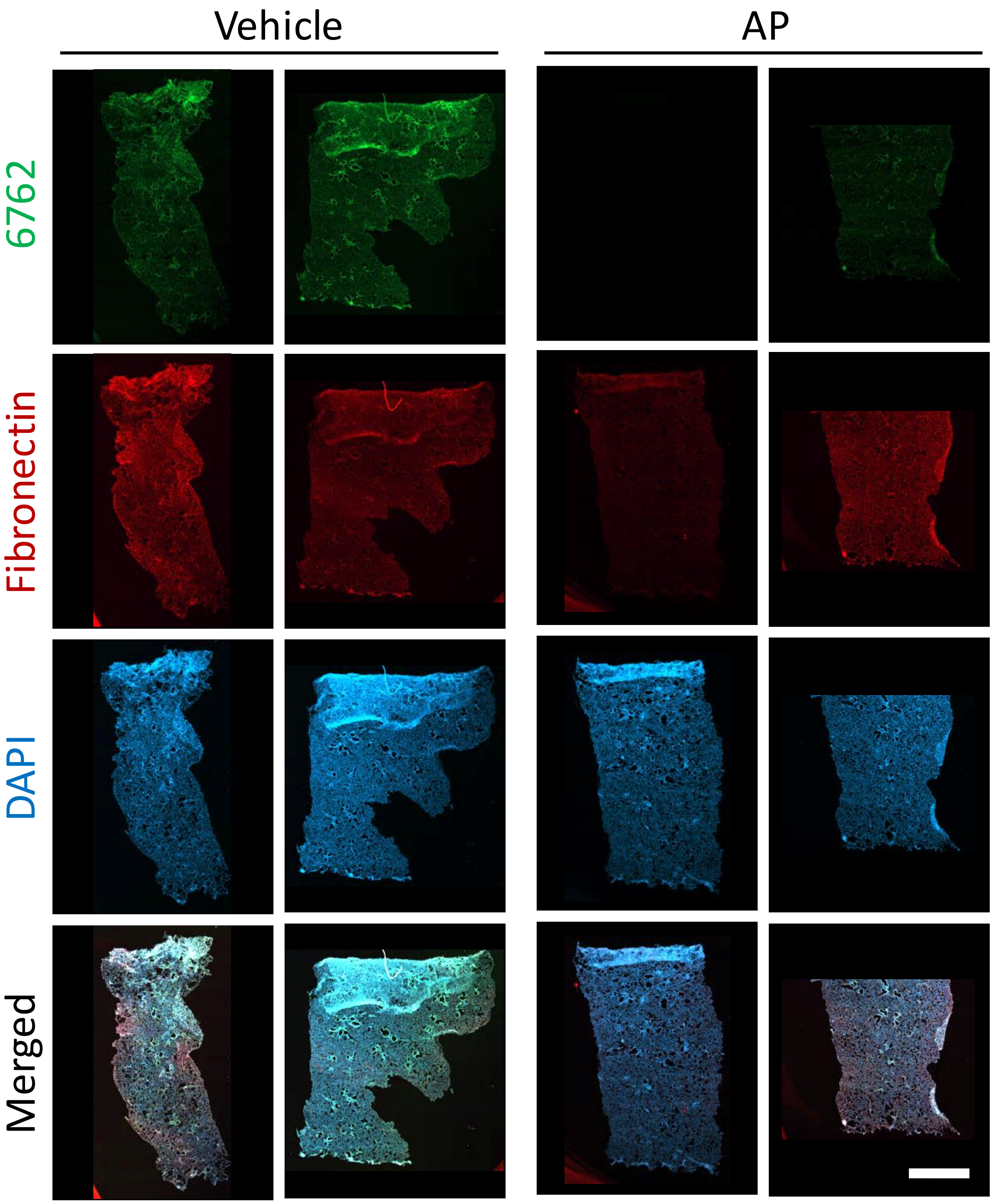

**Figure S17. Removal of p16-positive cells leads to reduced aptamer 6762 staining throughout the entire tissue section.** Aptamer 6762 is detected by fluorescein label (green). Fibronectin antibody is detected with AlexaFluor594 anti-rabbit secondary (red). Nuclei are stained with DAPI (blue). The images are tiled confocal images taken at 5× of the entire tissue sections. Scale bar (lower right panel) is 2 mm.

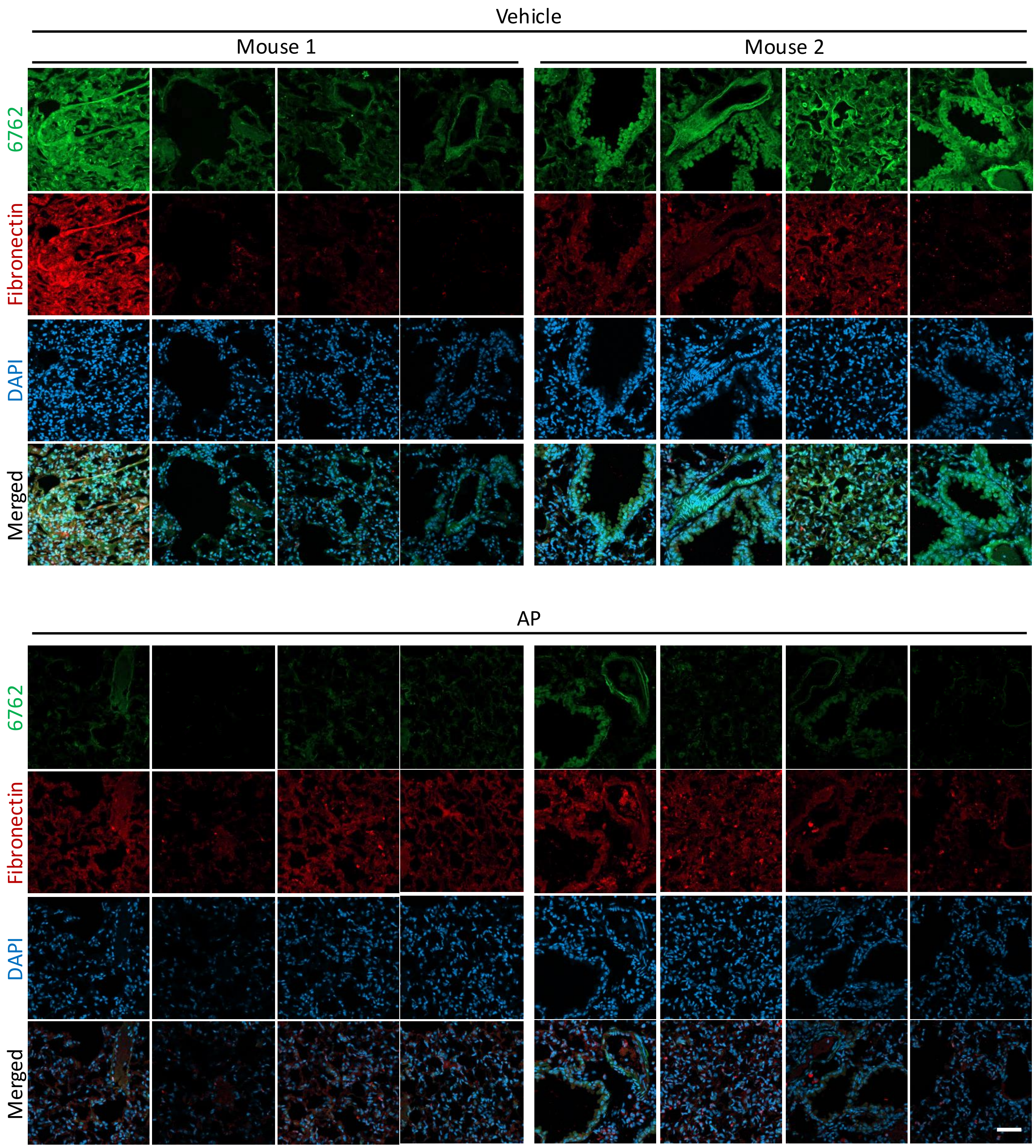

**Figure S18. Additional images show removal of p16-positive cells leads to reduced aptamer 6762 staining.** Aptamer 6762 is detected by fluorescein label (green). Fibronectin antibody is detected with AlexaFluor594 anti-rabbit secondary (red). Nuclei are stained with DAPI (blue). Scale bar (lower right panel) is 50 μm.

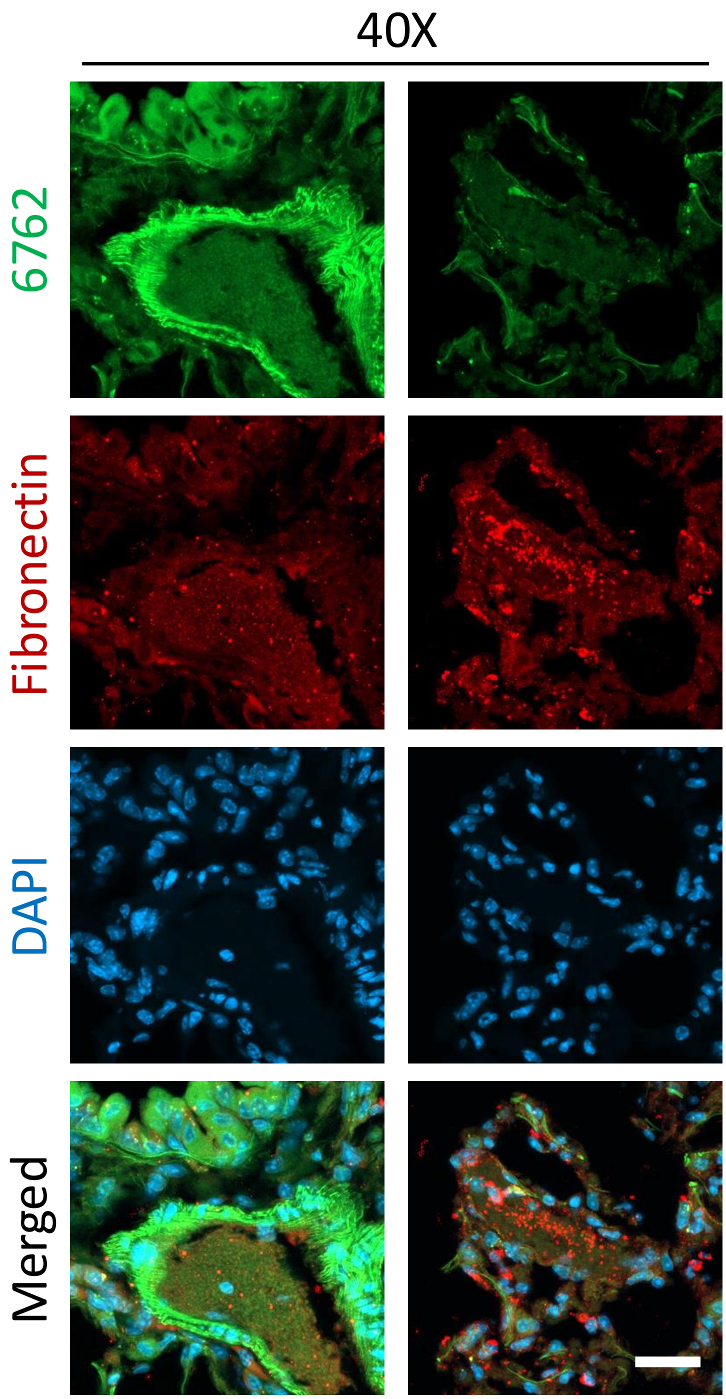

**Figure S19. Confocal imaging of mouse lung tissue with aptamers and anti-fibronectin antibody do not show complete overlap.** FAM-labeled aptamer 6762 (green) does not stain in an identical pattern as anti-fibronectin antibody detected by AlexaFluor594 labeled anti-rabbit secondary (red). Nuclei are stained with DAPI (blue). Scale bar (lower right panel) is 25 µm.

**Supplementary References**

1. Seluanov, A., Vaidya, A. and Gorbunova, V. (2010) Establishing primary adult fibroblast cultures from rodents. *J Vis Exp*.
2. Pearson, K., Doherty, C., Zhang, D., Becker, N.A. and Maher, L.J., 3rd. (2022) Optimized quantitative PCR analysis of random DNA aptamer libraries. *Anal Biochem*, **650**, 114712.
3. Edgar, R.C. (2010) Search and clustering orders of magnitude faster than BLAST. *Bioinformatics*, **26**, 2460-2461.
4. Shen, W., Le, S., Li, Y. and Hu, F. (2016) SeqKit: A Cross-Platform and Ultrafast Toolkit for FASTA/Q File Manipulation. *PLoS One*, **11**, e0163962.
5. Hoinka, J., Backofen, R. and Przytycka, T.M. (2018) AptaSUITE: A Full-Featured Bioinformatics Framework for the Comprehensive Analysis of Aptamers from HT-SELEX Experiments. *Mol Ther Nucleic Acids*, **11**, 515-517.
6. Bing, T., Shangguan, D. and Wang, Y. (2015) Facile Discovery of Cell-Surface Protein Targets of Cancer Cell Aptamers. *Mol Cell Proteomics*, **14**, 2692-2700.
7. Cox, J. and Mann, M. (2008) MaxQuant enables high peptide identification rates, individualized p.p.b.-range mass accuracies and proteome-wide protein quantification. *Nat Biotechnol*, **26**, 1367-1372.
8. Kalari, K.R., Nair, A.A., Bhavsar, J.D., O'Brien, D.R., Davila, J.I., Bockol, M.A., Nie, J., Tang, X., Baheti, S., Doughty, J.B. *et al.* (2014) MAP-RSeq: Mayo Analysis Pipeline for RNA sequencing. *BMC Bioinformatics*, **15**, 224.
9. Dobin, A., Davis, C.A., Schlesinger, F., Drenkow, J., Zaleski, C., Jha, S., Batut, P., Chaisson, M. and Gingeras, T.R. (2013) STAR: ultrafast universal RNA-seq aligner. *Bioinformatics*, **29**, 15-21.
10. Liao, Y., Smyth, G.K. and Shi, W. (2013) The Subread aligner: fast, accurate and scalable read mapping by seed-and-vote. *Nucleic Acids Res*, **41**, e108.
11. Robinson, M.D., McCarthy, D.J. and Smyth, G.K. (2010) edgeR: a Bioconductor package for differential expression analysis of digital gene expression data. *Bioinformatics*, **26**, 139-140.
12. Prieto, L.I., Sturmlechner, I., Graves, S.I., Zhang, C., Goplen, N.P., Yi, E.S., Sun, J., Li, H. and Baker, D.J. (2023) Senescent alveolar macrophages promote early-stage lung tumorigenesis. *Cancer Cell*, **41**, 1261-1275 e1266.
